# Conformations and sequence determinants in the lipid binding of an adhesive peptide derived from *Vibrio cholerae* biofilm

**DOI:** 10.1101/2025.07.14.664771

**Authors:** Xin Huang, Ramesh Prasad, Sarvagya Saluja, Yiyan Yang, Qi Yan, Sydney O. Shuster, Rich Olson, Chenxiang Lin, Caitlin M. Davis, Xiaofang Jiang, Huan-Xiang Zhou, Jing Yan

**Author notes:** These authors made equal contributions (Xin Huang, Ramesh Prasad). Correspondence (H.-X.Z.) and (J.Y.).

## Abstract

Surface adhesion is critical to the survival of pathogenic bacteria both in natural niches and during infections, often via forming matrix-embedded communities called biofilms. We previously identified a 57-amino acid peptide (Bap1-57aa) as a key contributor to biofilm adhesion of the pandemic pathogen *Vibrio cholerae* to various surfaces including lipid membranes. Here, we combine biophysical, computational, and genetic approaches to elucidate the molecular mechanism. A central aromatic-rich motif anchors the peptide to lipid bilayers while peripheral pseudo repeats enhance binding through avidity. Surprisingly, the core motif undergoes a lipid-induced conformational transition into a β-hairpin, enabling robust membrane insertion. Moreover, the biofilm-derived peptide, conserved in several other Vibrio species, can adhere to model host surfaces and is sensitive to membrane curvature. Our results provide molecular insight into biofilm adhesion and may lead to new strategies for targeted biofilm removal and the design of bioinspired underwater adhesives.

**Teaser:** A short peptide from *Vibrio cholerae* binds lipids using a unique β-hairpin motif and contributes to host colonization.

## Introduction

Membrane-interacting peptides are integral to a wide range of biological processes (*1*). These peptides influence membrane permeability, curvature, and organization, playing critical roles in cellular signaling, regulation, and transport (*2*). For instance, antimicrobial peptides kill bacteria by targeting and disrupting bacterial membranes or membrane crossing (*3–5*), while many pathogens secrete toxins or virulence factors that target host cell membranes to facilitate infection (*6*, *7*). Additionally, peptide-lipid interactions are implicated in neurodegenerative diseases such as Alzheimer’s disease, where amyloid β (Aβ) peptides aggregate on lipid membranes and compromise the neuronal membrane (*8*, *9*).

While peptide-lipid interactions are well-documented for certain motifs such as amphipathic helices and pore-forming peptides (*10–12*), the diversity of mechanisms and conformations by which peptides interact with lipid bilayers remains to be fully explored. Moreover, the biological functions of the lipid-peptide interactions remain poorly defined in many cases; in particular, much less understood are lipid-peptide interactions in the context of the bacteria-host interface, including their roles in colonization and host response. From an application perspective, such studies may inspire the design of biomimetic adhesives, which are critically needed in biomedical applications, especially for adhesion to wet surfaces.

Biofilms are surface-associated bacterial communities encased in an extracellular, polymeric matrix (*13–16*). Although many bacteria rely on this lifestyle for their survival in nature, biofilms are also widely found in infections and biofouling (*17*, *18*). In a prior study of biofilms formed by *Vibrio cholerae*, the causal agent of the pandemic cholera (*19*, *20*), we found that two partially redundant matrix proteins, Bap1 and RbmC, behave as double-sided tape to anchor *V. cholerae* biofilm clusters to host and environmental surfaces (*21*). Specifically, both proteins bind to the major biofilm matrix component <u>V</u>ibrio <u>p</u>oly<u>s</u>accharide (VPS) via a conserved β-propeller domain while containing a diverse set of surface binding functionalities in their environment-facing domains. Bap1 adheres to abiotic surfaces and lipid membranes via a 57 amino-acid loop (hitherto called Bap1-57aa) nested in a β-prism domain, whereas RbmC possesses domains targeting O-glycan-containing mucins and complex N-glycans prevalent in host cell-surface proteins (*10*). Together, Bap1 and RbmC play critical roles in enabling *V. cholerae* biofilms to attach to environmental and host cell surfaces, enhancing colonization and potentially pathogenicity (*22*, *23*).

In our previous study (*21*), we found that the chemically synthesized peptide with the Bap1-57aa sequence can bind to lipid membranes and various abiotic surfaces outside of the biofilm context. We further proposed that Bap1-57aa might be used as a generic and readily manipulatable underwater glue. However, the molecular mechanism underlying its adhesion, particularly to lipids, remains unresolved. Here, we employ an integrative approach combining biofilm assays, molecular dynamics (MD) simulations, bioinformatics, and conformational analyses to dissect how Bap1-57aa interacts with lipids to potentially allow *V. cholerae* to adhere to plasma membranes of epithelial cell surfaces during infection. These findings provide mechanistic insights into the collective adhesion of *V. cholerae* cells and establish Bap1-57aa as a valuable model for studying host-pathogen interactions involving biofilms.

## Results

### Molecular dynamics simulations reveal a core motif of Bap1-57aa for lipid binding

The Bap1-57aa sequence is enriched in aromatic and basic amino acids, arranged in four pseudo repeats with a consensus sequence WbpKpnmY, where b, p, n, and m denote basic (R, K, or H), polar (T, Q, or E), nonpolar (V or I), and mixed (P, A, or S) amino acids, respectively (Fig. 1a and Fig. S1). The linker between the two inner pseudo repeats, WFFG, is also enriched in aromatic amino acids. To gain insight into the interaction of Bap1-57aa with lipids, we performed all-atom MD simulations of Bap1-57aa interacting with a lipid bilayer (POPC:POPS:PIP_2_ at 75:20:5 molar ratios, modeling the plasma membrane (*24*)). Without prior knowledge of possible conformations of the peptide, we started from extended conformations generated by the TraDES method (*25*), which produces conformations of intrinsically disordered proteins (IDPs) from their sequences, as done in our previous simulations of IDP-membrane systems (*26*). After being placed near the bilayer surface, Bap1-57aa first attaches to the bilayer via basic residues in the pseudo repeats. Subsequently, it spontaneously inserts into the bilayer via the central linker in three of eight replicate simulations during the first 100-350 ns (Fig. S2a, b). This set of simulations is referred to as IDP-wor (the last term for without restraint).

**Figure 1.**
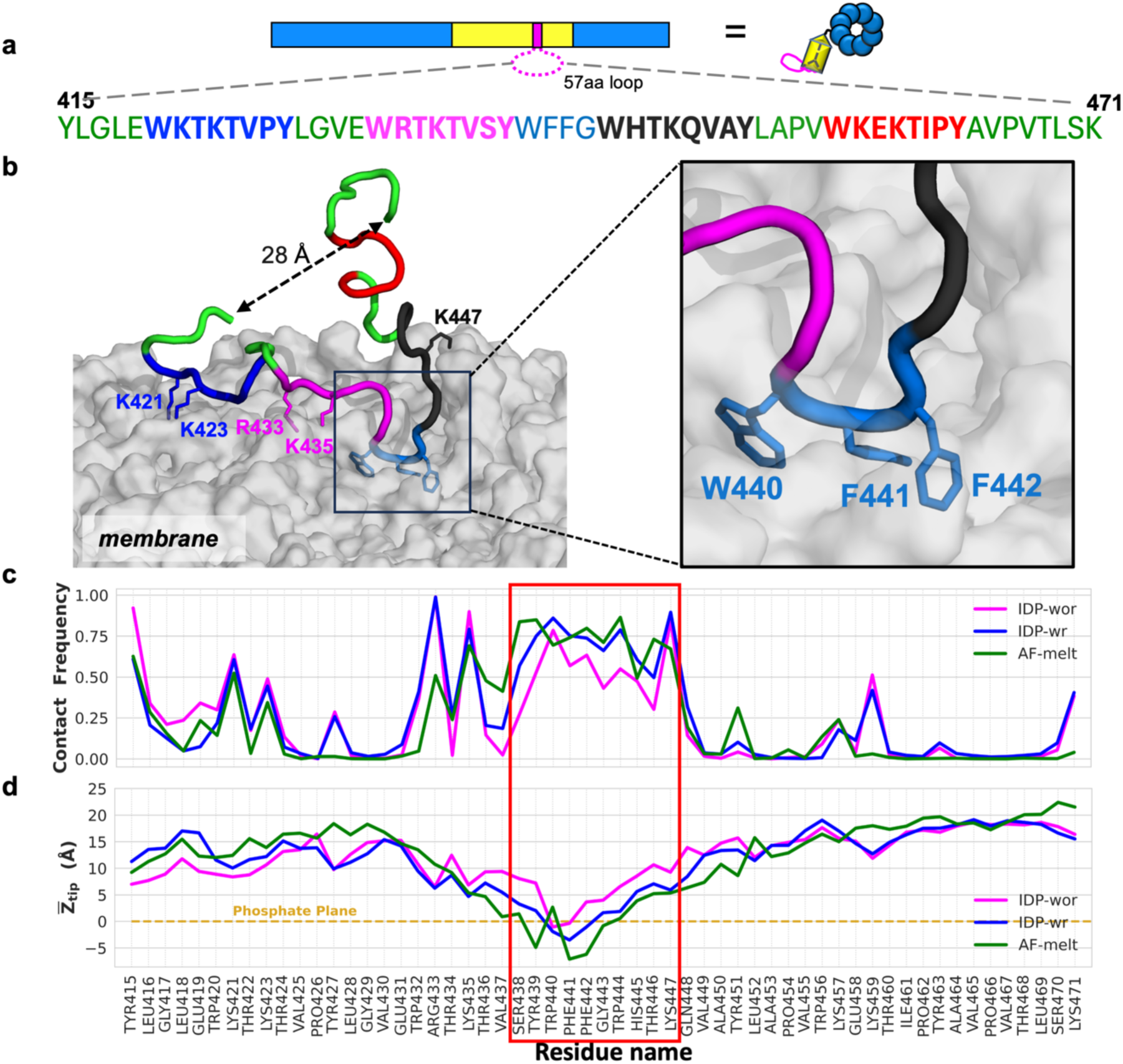
A core motif in the biofilm-derived peptide is identified as key for lipid binding through MD simulations. (**a**) Schematic of Bap1 structure and sequence of the Bap1-derived peptide (residues 415-471, abbreviated as Bap1-57aa). Bap1-57aa corresponds to a loop in the biofilm-specific adhesion molecule Bap1. Four pseudo repeats in the Bap1-57aa sequence are shown in blue, magenta, black, and red. The core hydrophobic motif (WFFG) are shown in cyan and the rest of the residues are shown in green. (**b**) Snapshot from a simulation of Bap1-57aa-membrane binding, started from a disordered conformation for the peptide with the Cα atoms of the N-and C-terminal residues restrained at 28 Å. The middle linker W_440_FFG_443_ inserts into the membrane (zoomed view). (**c**) Membrane contact frequencies from the simulations without N-C distance restraint (IDP-wor), with N-C distance restraint (IDP-wr), and in a conformation melted from the AlphaFold structure (AF-melt). (**d**) Mean Z_tip_ values from the IDP-wor, IDP-wr, and AF-melt simulations, respectively.

In the full-length Bap1, the 57aa peptide is a loop nested in a β-prism domain, with the end-to-end distance restrained. To mimic this situation, in a second set of 12 simulations termed IDP-wr (with restraint), we restrained the end-to-end Cα-Cα distance of Bap1-57aa to 28 Å, which is the mean value in the IDP-wor simulations during the first 450 ns (Fig. S2c). A representative snapshot from the IDP-wr simulations is presented in Fig. 1b, showing robust bilayer insertion of the central linker along with lipid binding of basic residues in the pseudo repeats. In a third set of four simulations (referred to as AF-melt), we started with a partially melted conformation (via simulations at 500 K) of a structure predicted by AlphaFold (*27*).

The three sets of simulations show similar membrane-binding features. For example, the membrane contact frequencies of individual residues have similar patterns, with the central segment, S_438_YWFFGWHTK_447_ (hereafter referred to as the “core motif”), exhibiting the highest membrane contact frequencies (Fig. 1c). The peripheral residues, especially basic residues K421, K423, R433, K435, and K459, also have significant membrane contact frequencies. As a complementary measure, we calculated Z_tip_, the Z coordinate (along the membrane normal) of the sidechain tip heavy atom of each residue relative to the proximal phosphorus plane (*28*). The mean Z_tip_ values of the middle linker are at or below 0, strongly indicative of membrane insertion, in all three sets of simulations (Fig. 1d).

### Fluorescence-based lipid adsorption assays confirm the importance of the core motif and avidity of the pseudo repeats

Inspired by the MD simulation results, we set out to test the central hypothesis that the aromatic core motif and the peripheral pseudo-repeats contribute jointly to the membrane interaction of Bap1-57aa. To systematically and quantitatively study Bap1-57aa-lipid interactions, we chemically synthesized the Bap1-57aa peptide with an N-terminal fluorescein isothiocyanate (FITC) label, and developed a protocol to visualize and quantify its spontaneous adsorption onto lipid-coated microbeads using fluorescence microscopy (Fig. 2a) (*21*, *29*). In brief, supported lipid bilayers were formed on 5 μm silica beads by incubating them with small unilamellar vesicles (SUVs) containing Rhodamine-labeled lipids. Subsequently, adsorption curves were generated by measuring excess fluorescence signals on the surface of the lipid-coated beads relative to the solution signals at a series of peptide concentrations. The fluorescence signal on the beads is assumed to be proportional to the number of peptide molecules bound to the surface-anchored lipids.

**Figure 2.**
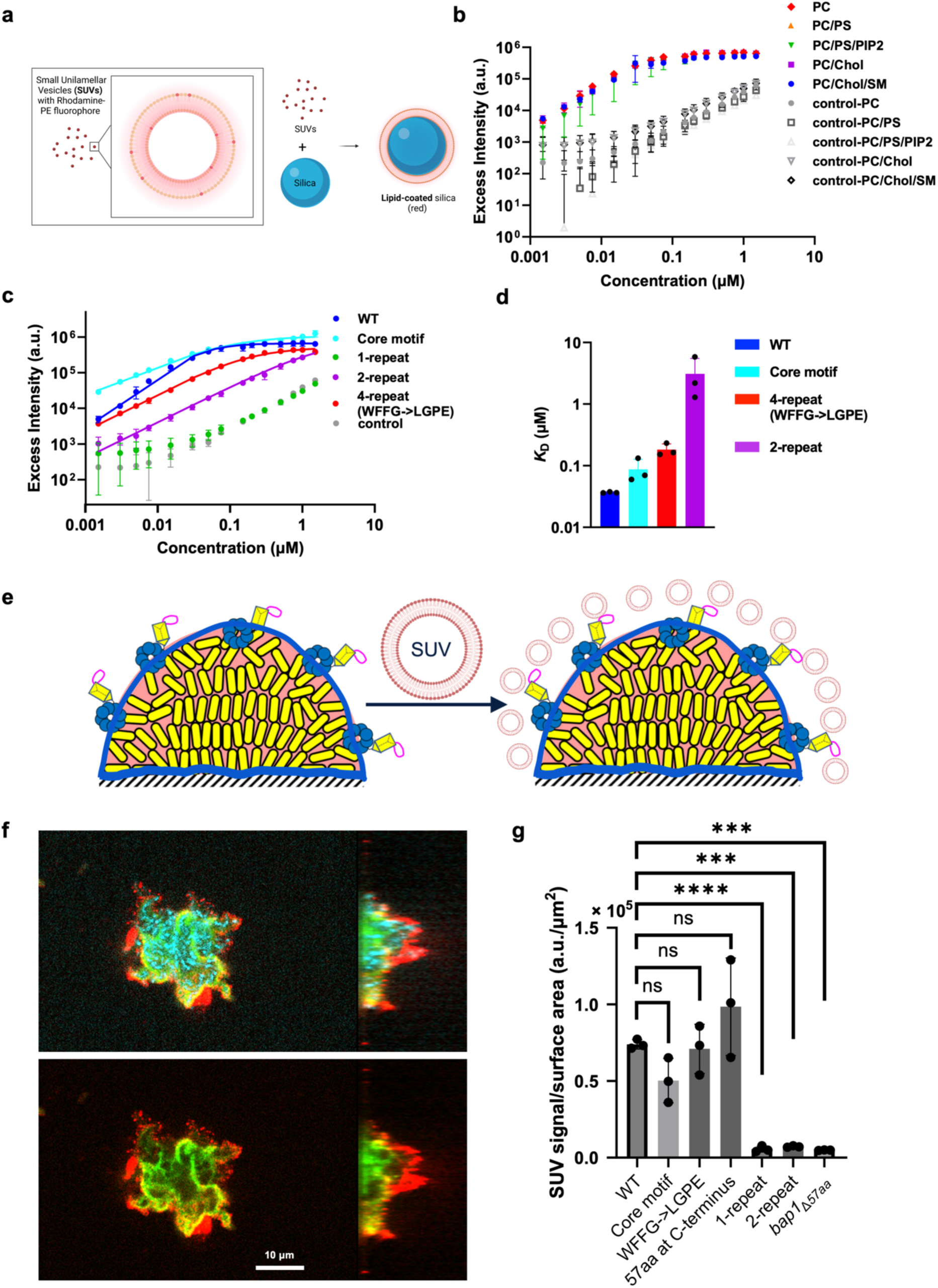
Experimental evidence for the importance of the core motif and avidity of the pseudo repeats in lipid binding. (**a**) Schematic of the fluorescence-based bead adsorption assay. (**b**) Robust lipid binding of Bap1-57aa with respect to lipid composition. Excess fluorescence signals on the bead surface relative to the solution signals are plotted against initial peptide concentration, using beads coated with various lipids. Dextran (4 kDa) conjugated to FITC was used as a negative control. DOPC = 1,2-dioleoyl-sn-glycero-3-phosphocholine. DOPS = 1,2-dioctadecenoyl-sn-glycero-3-phosphoserine. PIP_2_ = phosphatidylinositol 4,5-bisphosphate. Chol = cholesterol. SM = sphingomyelin. (**c**) Adsorption curves with Hill equation fitting for Bap1-57aa and sequence variants on DOPC-coated beads. (**d**) Dissociation constants derived from fitting adsorption curves to a Hill equation, for peptides with different sequences. (**e**) Schematic of the biofilm-based adsorption assay. Rh-PE labeled SUVs were added to a confluent layer of biofilm grown from *V. cholerae* cells constitutively expressing mNeonGreen to assess the ability of the biofilm to capture SUVs. (**f**) Cross-sectional images of a biofilm from cells constitutively expressing SCFP3A (cyan); Bap1 was tagged with 3×FLAG and labeled with anti-FLAG antibody conjugated to FITC (green). Rh-PE labeled SUVs (red) bind to the periphery of the cell cluster. (**g**) Quantification of SUV capture by biofilms formed by various *V. cholerae* mutants using the total SUV signal intensity normalized by the biofilm surface area. a.u. stands for arbitrary unit. All data represent mean ± SD (*n* = 3 biological replicates). Statistical analyses were performed using two-tailed *t*-tests with Welch’s correction. ns, not significant; ***, *p* < 0.001; ****, *p* < 0.0001. *p* values from left to right: 0.1006, 0.7815, 0.3136, <0.0001, 0.0005, and 0.0006.

Given the complexity and diversity of cell membranes, which contain various lipids including cholesterol (*30*), we tested the wild-type (WT) Bap1-57aa peptide with various lipid compositions. In general, the adsorption curves largely overlap and show a rapid increase up to 0.05 µM peptide concentration and a plateau thereafter (Fig. 2b). Although the fluorescence signals in the negative control (FITC-labeled dextran) also increase with increasing concentrations, possibly due to weak nonspecific binding, the signals are two orders of magnitude lower than those for WT Bap1-57aa regardless of the lipid composition. These observations demonstrate that the binding between the Bap1-57aa peptide and lipid bilayers is robust with respect to variations in lipid composition.

After establishing the binding assay, we sought to elucidate the lipid binding mechanism of Bap1-57aa by designing and testing a series of peptide variants (Fig. S1b, Fig. 2c). Indeed, we found that the core motif displays strong adsorption despite its much shorter length, consistent with the MD simulation results. In contrast, a representative 1-repeat unit (WKTKTVPY) does not show significant lipid binding, as its adsorption curve is close to the negative control. Interestingly, as the number of the repeating units increases, the ability to adsorb onto lipids improves. For example, a 2-repeat peptide shows a significant upward shift in the binding curve. We also designed a variant with a central linker lacking the aromatic residues (WFFG→LGPE), to isolate the contributions of the four pseudo repeats; this variant shows even stronger adsorption to lipids than the 2-repeat peptide. Fitting the adsorption curves to the classical Hill binding model yielded a low dissociation constant (*K*_D_) of 37±1 nM for the WT peptide, indicating a high binding affinity (Fig. 2d). By comparison, the binding affinities of the core motif and the WFFG→LGPE variant are lowered by 2.4 and 5.0-fold, respectively, compared to the WT peptide. Moreover, the affinity of the 2-repeat variant is further reduced from the 4-repeat variant (WFFG→LGPE) by 17-fold and the 1-repeat unit has no measurable binding, suggesting an avidity effect (*1*, *26*, *31*), i.e., each pseudo repeat may bind lipids weakly but their simultaneous presence strengthens the binding significantly. Overall, these results suggest synergistic effects of the aromatic core motif and the peripheral pseudo repeats, resulting in the strong adsorption of Bap1-57aa to lipid-coated surfaces.

### Bap1-57aa plays a key role in the binding of *V. cholerae* biofilms to host membranes

Next, we tested the relevance of the *in vitro* observations in the native context, *V. cholerae* biofilms. To do so, we first generated *V. cholerae* mutants containing various *bap1* constructs. In parallel, we developed another binding assay in which a mature biofilm of *V. cholerae* cells constitutively expressing mNeonGreen was grown on glass, followed by the introduction of Rhodamine-labeled lipid SUVs into the culture (Fig. 2e, f). After vigorous washing, we quantified the remaining Rhodamine signals, normalized by the total biofilm surface area, as a measure of the interaction strength between the biofilm surface and the SUVs (Fig. 2g). Through hydrophobicity measurements (Fig. S3) (*32*), we inferred that the secreted Bap1 molecules adopt a configuration with the Bap1-57aa loop facing the environment and the β-propeller facing towards and binding to the biofilm matrix. We hypothesize that this configuration allows Bap1 to interact with and capture exogenously added SUVs. In this assay, we used a lipid composition of DOPC-sphingomyelin-cholesterol since it resembles the composition of the outer leaflet of the plasma membrane in mammalian cells (*30*, *33*, *34*), which is most likely encountered by *V. cholerae* cells during an infection after mucosal penetration.

To demonstrate the validity of the assay, we first verified that WT *bap1* biofilms show strong SUV capturing abilities while the negative control with the Bap1-57aa loop deleted (*bap1*_Δ*57aa*_) did not (Fig. 2g, WT vs. *bap1*_Δ*57aa*_). Next, we generated several Bap1-57aa variants in *V. cholerae* and tested the ability of the corresponding biofilms to capture SUVs. Notably, replacing the entire Bap1-57aa sequence with the 10-aa core motif does not affect the SUV capturing ability, highlighting its critical role (Fig. 2g, WT vs. Core motif). In contrast, the 1-repeat peptide (WKTKTVPY), despite having a similar length to the core motif (8 aa vs. 10 aa), is defective in SUV binding (Fig. 2g, 1-repeat). Adding a second repeat did not improve SUV capture, but the 4-repeat variant (WFFG→LGPE) exhibited SUV capture levels similar to WT, reaffirming the avidity effect in the peripheral pseudo repeats (Fig. 2g, 2-repeat vs. WFFG→LGPE). The loss of function in the defective mutants is unlikely due to changes in the secretion levels of the mutant proteins, as shown by the quantification of the amount of Bap1 molecules in biofilms using *in situ* immunostaining (Fig. S4b, c) (*21*, *35*, *36*). We were initially concerned that the cap of the β-prism domain adjacent to the Bap1-57aa loop, which has several solvent-exposed lysine and tryptophan residues (*37*), might complicate the interpretation of the data and mask functional changes in the WFFG→LGPE mutant. To address this confounding factor, we additionally constructed mutants in a *bap1*_Δ*β-prism*_ background to isolate the effects of the Bap1-57aa loop. In this cleaner context, the WFFG→LGPE mutant showed significantly reduced SUV capture compared to WT Bap1-57aa, suggesting the indispensable role of the core motif (Fig. S4d, e).

To bridge the gap between the biofilm-based assay, where Bap1-57aa is a loop nested in a well-folded domain, and the *in vitro* assays in which the peptide lacks end-to-end restraints, we constructed a mutant in which the 57aa sequence was repositioned at Bap1’s C-terminus, thus releasing the end-to-end restraints on the peptide. Interestingly, this mutant maintains a WT-level lipid-binding ability (Fig. 2g, Fig. S4d), suggesting that the Bap1-57aa peptide can function independently of its original loop context regarding lipid binding – consistent with the *in vitro* results with chemically synthesized peptides that have no such conformational restraints.

### Bap1-57aa core motif binds lipid membranes in a β-hairpin conformation

We next used several biophysical methods to probe the molecular details of the 57aa-lipid interaction. Because MD simulations predict that aromatic sidechains in the core motif may insert into the membrane and provide a stable anchor (Fig. 1), we started by focusing on this sequence. We first used fluorescence spectroscopy to examine the environment of the tryptophan residues within the core motif in the presence and absence of SUVs (DOPC/DOPS 3:1). At a fixed peptide concentration, increasing the amount of SUVs significantly enhances tryptophan fluorescence intensity (Fig. 3a). The emission peak also exhibits a noticeable blue shift upon addition of lipids. We reason that, in the absence of lipids, the tryptophan residues are solvent-exposed, whereas in the presence of lipids, the tryptophan residues insert into the hydrophobic core of the lipid bilayer (*38*). With more SUVs available, a higher fraction of peptide molecules binds to the bilayer, likely inserting into the membrane, resulting in the observed fluorescence enhancement and spectral shift. The full-length Bap1-57aa peptide shows a peak at 340 nm even in the absence of lipids, which is already significantly blue-shifted relative to free tryptophan in solution. This suggests that the tryptophan residues in the WT Bap1-57aa peptide may already be in a constrained conformation, e.g., due to burial in a collapsed chain in solution. As a consequence, a minimal blue shift is observed in the emission peak when SUVs are present, though the peak intensity shows a similar increase (Fig. 3b). In comparison, the WFFG→LGPE variant peptide shows increased fluorescence intensity in the presence of SUVs confirming lipid binding, but the peak wavelength in the absence of SUVs is reminiscent of free tryptophan in solution and shows only a very modest blue shift in the presence of SUVs (Fig. S5). The latter observation suggests that peripheral tryptophans are not as deeply buried as the core tryptophans, consistent with MD simulations (Fig. 1d).

**Figure 3.**
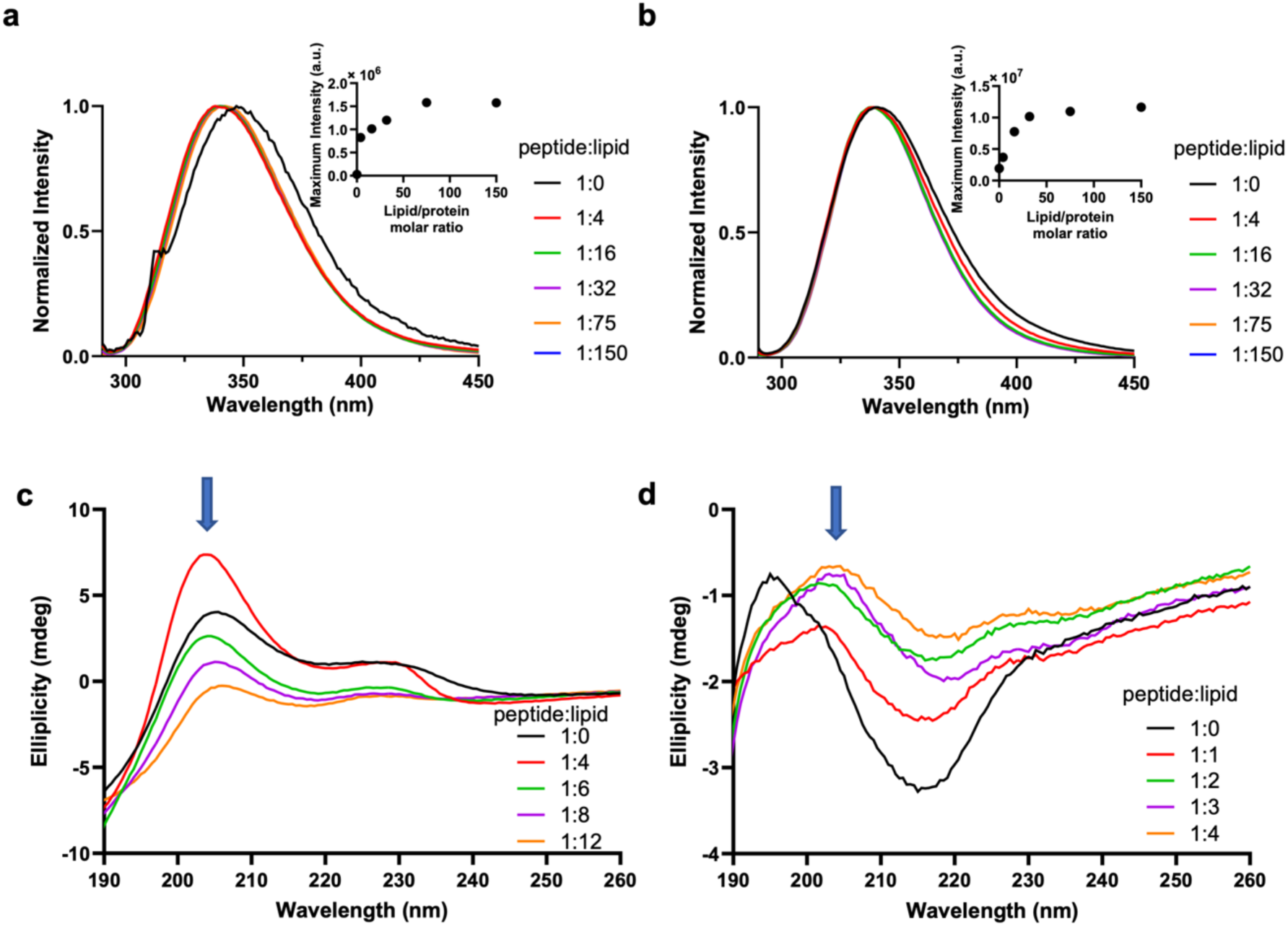
The core motif binds lipid membranes in a β-hairpin conformation. (**a-b**) Normalized tryptophan fluorescence spectra of (**a**) 3 μM core motif or (**b**) 3 μM Bap1-57aa mixed with 3:1 DOPC/DOPS SUV at molar ratios between 1:0 and 1:150 in 10 mM Tris buffer pH 7.4 and 150 mM NaCl. Insets: Maximum tryptophan fluorescence intensity of the corresponding peptide plotted over a series of peptide:lipid molar ratios. (**c**) CD spectra of 50 μM core motif mixed with 3:1 DOPC/DOPS SUV at peptide:lipid molar ratios between 1:0 and 1:12 in 2 mM Tris buffer pH 7.4 and 5 mM NaCl at 20 °C. The characteristic peak at 205 nm indicates a β-turn structure. (**d**) CD spectra of 25 μM Bap1-57aa peptide mixed with 3:1 DOPC/DOPS SUV at molar ratios between 1:0 and 1:4 in 2 mM Tris buffer pH 7.4 and 5 mM NaCl at 20 °C. a.u. stands for arbitrary unit.

To generate further mechanistic insights, we investigated whether the Bap1-57aa peptide exhibits any secondary structures when interacting with lipids. First, we collected far-UV circular dichroism (CD) spectra of the core motif peptide (50 μM) mixed with SUVs (3:1 DOPC/DOPS) at varying molar ratios (Fig. 3c). SUVs are optically transparent in CD, allowing us to test the secondary structures of the peptide when binding to lipids. At high lipid concentrations, the amplitude of the CD spectra decreases, which is a common optical artifact observed in studies of SUV-protein interactions (*39*).

Notably, in the absence of the lipids, the CD spectrum of the core motif shows a positive band at ∼205 nm, characteristic of a type II β-turn (*40*, *41*). Type II β-turns have been shown to exhibit CD spectra with a strong positive band between 200 and 210 nm and a shallow negative band between 220 and 225 nm, in contrast to the CD spectra of classical β-sheets with positive bands at 195 nm and negative bands at 218 nm (*42*). Additionally, we observed a peak at ∼230 nm in the CD spectra, attributable to tryptophan coupling effects and restraints in tryptophan rotation (*43*, *44*); this is again consistent with a β-turn conformation of the core motif in which the two tryptophan residues are brought into close proximity. Thus, the CD data suggest that the core motif peptide may adopt a β-hairpin conformation with a turn in the middle, which also agrees with AlphaFold predictions (Fig. S1c) (*27*).

Next, to test if the β-hairpin conformation is preserved in the full Bap1-57aa sequence, we collected far-UV CD spectra of the Bap1-57aa peptide with and without SUVs (Fig. 3d). The CD spectrum of Bap1-57aa in the absence of SUVs exhibits a positive band at 195 nm and a strong negative band between 210 and 220 nm, indicating the formation of classical β-sheets. However, with increasing levels of SUVs, both bands recede and the characteristic type II β-turn 205 nm peak emerges (originally a shoulder in the lipid-free CD spectrum), indicating that the Bap1-57aa peptide gains a β-hairpin conformation when interacting with lipid membranes. It appears that lipid binding reinforces the intrinsic tendency of the core motif to form a β-hairpin conformation, promoting its insertion and anchoring the entire peptide to the membrane.

### MD simulations of membrane binding of Bap1-57aa in a β-hairpin conformation

Motivated by the CD data implicating a β-hairpin conformation, we carried out a new set of MD simulations of Bap1-57aa binding with membranes, now with the core motif adopting a β-hairpin conformation, with the FG residues of the central linker WFFG at the *i* + 1 and *i* + 2 positions of the turn (Fig. 4a). The MD simulations in this new conformation was performed both in solution and at the membrane surface. Interestingly, the β-hairpin conformation is stable only when interacting with lipids (Fig. 4b). In solution, the β-hairpin melts; instead, parallel or antiparallel β-sheets are formed between two strands away from the core motif (Fig. 4c). The contrasting behavior in solution and at the membrane surface is consistent with the conformational changes upon lipid binding suggested by the CD spectra (Fig. 3d).

**Figure 4.**
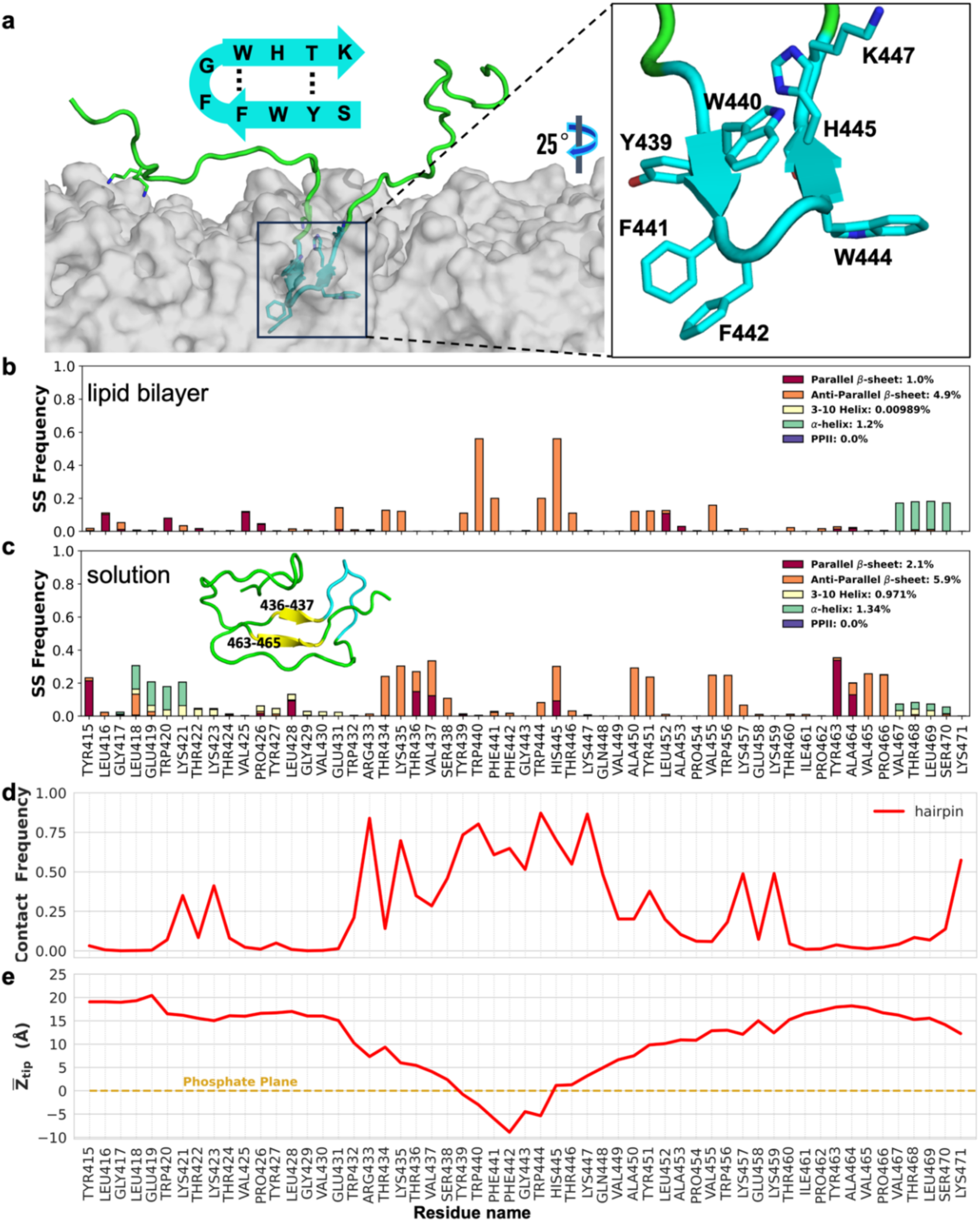
Simulation of the Bap1-57aa peptide interacting with membranes in a β-hairpin conformation. (**a**) A snapshot showing the core motif (S_438_YWFFGWHTK_447_; cyan) inserting into the lipid bilayer as a β-hairpin. Inset: schematic of the β-hairpin model for the core motif. (**b-c**) Frequency of secondary structures (SS) for each residue in simulations starting from a β-hairpin conformation, (**b**) at the lipid bilayer surface or (**c**) in solution. Inset in (c) shows a representative snapshot of Bap1-57aa in solution. (**d**) Contact frequency and (**e**) Z_tip_ distance from simulations starting from the β-hairpin conformation. Eight independent simulations were performed for 1.1 µs each; average properties are plotted.

Overall, the membrane-binding features in the set of simulations with the β-hairpin conformation (Fig. 4d, e) are similar to those in the preceding three sets of simulations (Fig. 1c, d). However, closer examination shows that the β-hairpin conformation allows deeper insertion of the hydrophobic residues at the tip of the β-hairpin (Fig. 4e). In particular, the two phenylalanine side chains (F_441_F_442_) are deeply buried in the hydrophobic core of the lipid bilayer in this configuration. Moreover, the tyrosine and histidine residues (Y_439_ and H_445_) sit at the phosphate plane, a preferred location for aromatic residues capable of forming hydrogen bonds (*45*, *46*). In contrast, peripheral aromatic residues show little propensity for membrane insertion, consistent with the contrasting behaviors in fluorescence peak shift between WT and WFFG→LGPE peptides (Fig. 3b, Fig. S5).

### Bap1-57aa binds to host cell surfaces

To demonstrate the relevance of our findings in the host context, we transitioned from the model lipid bilayers to the more complex, physiologically relevant cell surfaces. To this end, we stained Caco-2 cells (as a model for intestinal epithelial cells) with FITC-labeled peptides, along with a DNA stain (DAPI) and an actin stain (AlexaFluor 647 phalloidin). Figure 5 shows that WT, core motif, and WFFG→LGPE peptides stained the entire cell surface of Caco-2 cells, while the 1-repeat and 2-repeat peptides exhibited weak signals similar to the negative control (FITC-dextran). These results closely parallel those in our *in vitro* binding assays.

**Figure 5.**
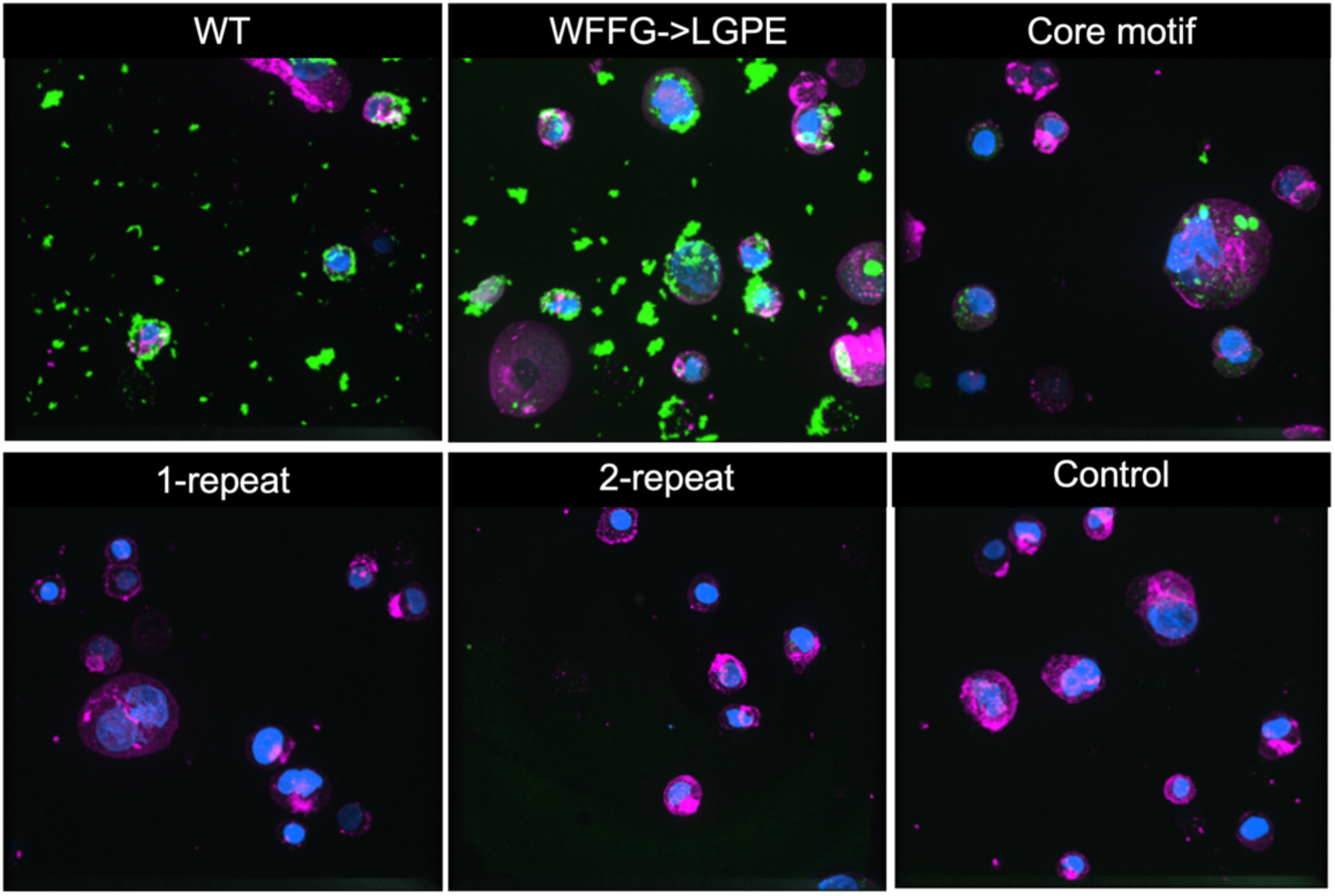
Bap1-57aa binds to cell surfaces. Shown are 3D rendering of confocal images of Caco-2 cells stained with 300 nM of DAPI (blue) for nuclei, 110 nM of AlexaFluor 647 phalloidin (magenta) for actin, and 1.5 µM FITC-labeled peptides (green). The total size of each image is 220 × 220 × 28 µm.

### Membrane binding of Bap1-57aa is curvature sensitive

The surfaces of intestinal epithelial cells are rich in membrane-bound protrusions called microvilli that have diameters on the order of tens to hundreds of nanometers (*47*). Therefore, we were curious to what extent the Bap1-57aa-lipid interaction is sensitive to local membrane curvature. To test whether Bap1-57aa binds membrane in a curvature-dependent manner, we prepared SUVs by sonication (mean diameter <40 nm) to mimic highly curved membrane features such as microvilli, and separately, large vesicles (LVs) by extrusion (mean diameter >150 nm) for comparison. We then measured the peptide-membrane binding using a co-flotation assay (Fig. 6a) (*48*, *49*), which separates a mixture of FITC-labeled peptides and Cy5-labeled vesicles in a sucrose gradient by the buoyant density of each species. After centrifugation, two major SUV populations were observed, one at the top of the gradient (fraction 1 or F1) and the other in the middle (F3). SDS-PAGE showed that the F1 fraction was mostly peptide-free (i.e., little FITC signal), while the F3 fraction contained a high level of the peptide (Fig. 6b). Negative stain transmission electron microscopy (TEM) showed that the peptide-free vesicles in F1 (61.8±23.7 nm, *n* = 73) were significantly larger than the peptide-bound vesicles in F3 (32.3±13.2 nm, *n* = 89), suggesting that Bap1-57aa selectively bound vesicles with higher membrane curvatures.

**Figure 6.**
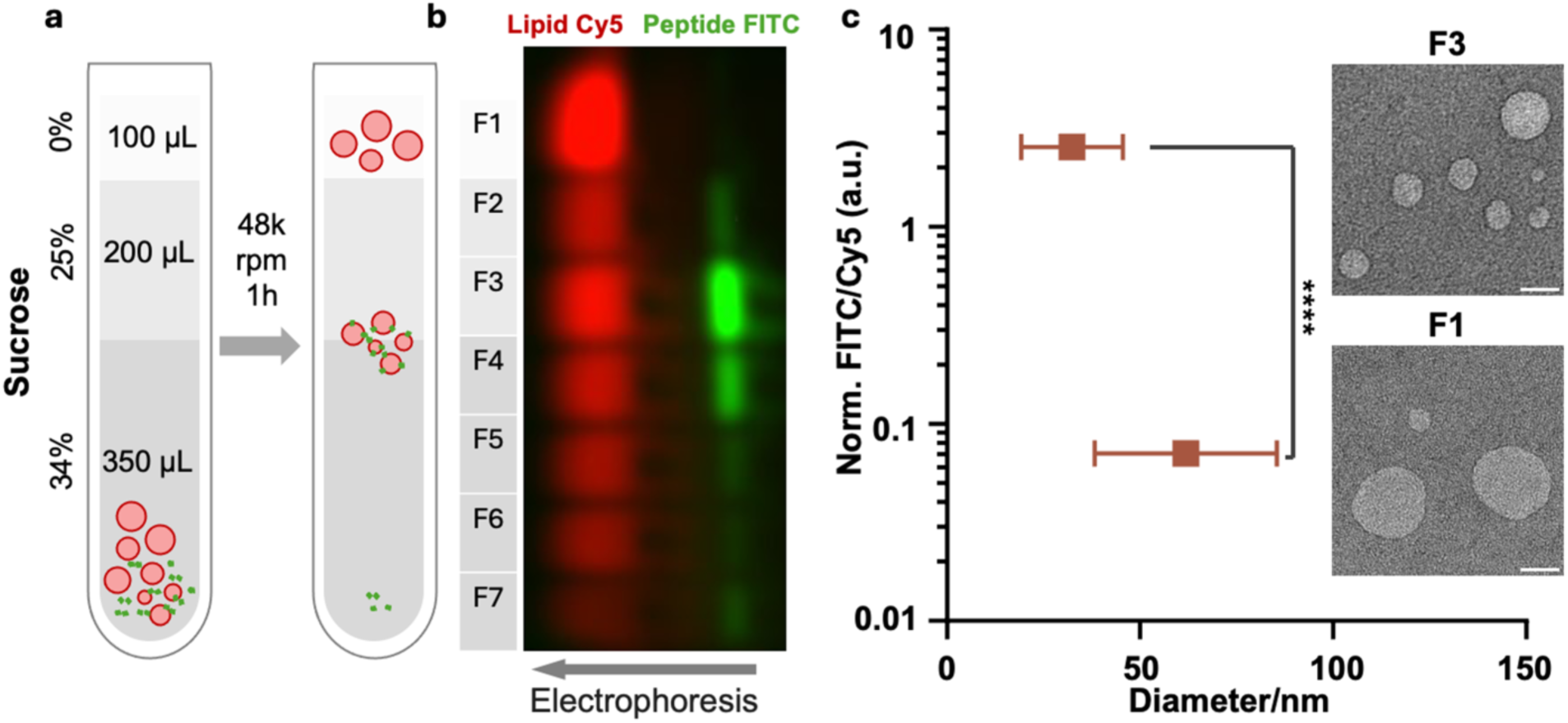
Bap1-57aa binding to SUVs analyzed by a flotation assay. (**a**) Schematic of the floatation assay. A mixture of SUVs (50 μM total lipid, containing 0.8% Cy5-DOPE) and FITC-labeled Bap1-57aa peptide (1 μM) are layered at the bottom of a sucrose density gradient. Centrifugation then separates peptide-free vesicles (top), peptide-bound vesicles (middle), and free peptides (bottom). (**b**) SDS-PAGE analysis of 7 fractions (100 μL each for F1–F6, 50 μL for F7) recovered from the post-centrifugation gradient. Pseudo-colors: Cy5, red; FITC, green. (**c**) FITC to Cy5 fluorescence ratio [FITC/Cy5, measured from the gel image in (**b**) and normalized (norm.) to ∑_*F1*_^*F7*^(FITC) / ∑_*F1*_^*F7*^(Cy5)], plotted as a function of mean diameter of vesicles (measured by negative-!# !# stain TEM) for fractions 1 and 3. Error bars show standard deviations, *n* = 73 (F1) and 89 (F3), ****: *p* < 0.0001 in Welch’s *t*-test. Scale bars for electron micrographs: 50 nm. a.u. stands for arbitrary unit. The experiments were repeated twice with similar results (see Fig. S6).

We further compared membrane binding of the peptide to SUVs and LVs by quantifying the flotation assay (Fig. S6). We found that in major factions containing peptide-coated vesicles, the peptide:lipid ratio on SUVs was about 2× higher than LVs. The peptide-bound LVs floated higher in the sucrose gradient than SUVs, consistent with LVs’ larger volume and more sparse peptide coating. We note that these results should be interpreted qualitatively, with the following caveats. First, the negative-stain TEM tends to overestimate vesicle size due to staining and drying. Second, a portion of vesicles generated by extrusion are multilamellar, meaning that part of the LV membranes may not be accessible to peptides. Nevertheless, our results clearly show a general preference of Bap1-57aa for high membrane curvature.

### Bap1-57aa is distributed across multiple Vibrio species with conserved sequences

Finally, to complement simulations and experiments and to put our findings into an evolutionary perspective, we performed bioinformatic analyses of Bap1-57aa. We first examined the presence of Bap1-57aa in all Vibrio proteins across the Vibrio genus identified from 6,121 genomes in the Genome Taxonomy Database (GTDB) r214 (*50*). This sequence was found exclusively in Bap1 proteins, highlighting its distinctive role. We then constructed a phylogenetic tree for all Bap1-encoded nucleotide sequences and mapped the presence or absence of the 57aa loop in various genomes (Fig. 7a). The 57aa loop is present among Bap1 proteins from seven *Vibrio* species, including *V. anguillarum*, *V. ordalii*, *V. metoecus*, and *V. cholerae*, with the majority (63%) originating from *V. cholerae*. Notably, within the Bap1 proteins of *V. cholerae* species, we identified a lineage lacking the 57aa loop (Fig. 7a). Note that the loop-less Bap1-encoded gene is a duplicate adjacent to the standard Bap1-encoded gene in the *V. cholerae* genome (*51*); it appears that after duplication, the loop is lost in some *V. cholerae* strains but retained in others.

**Figure 7.**
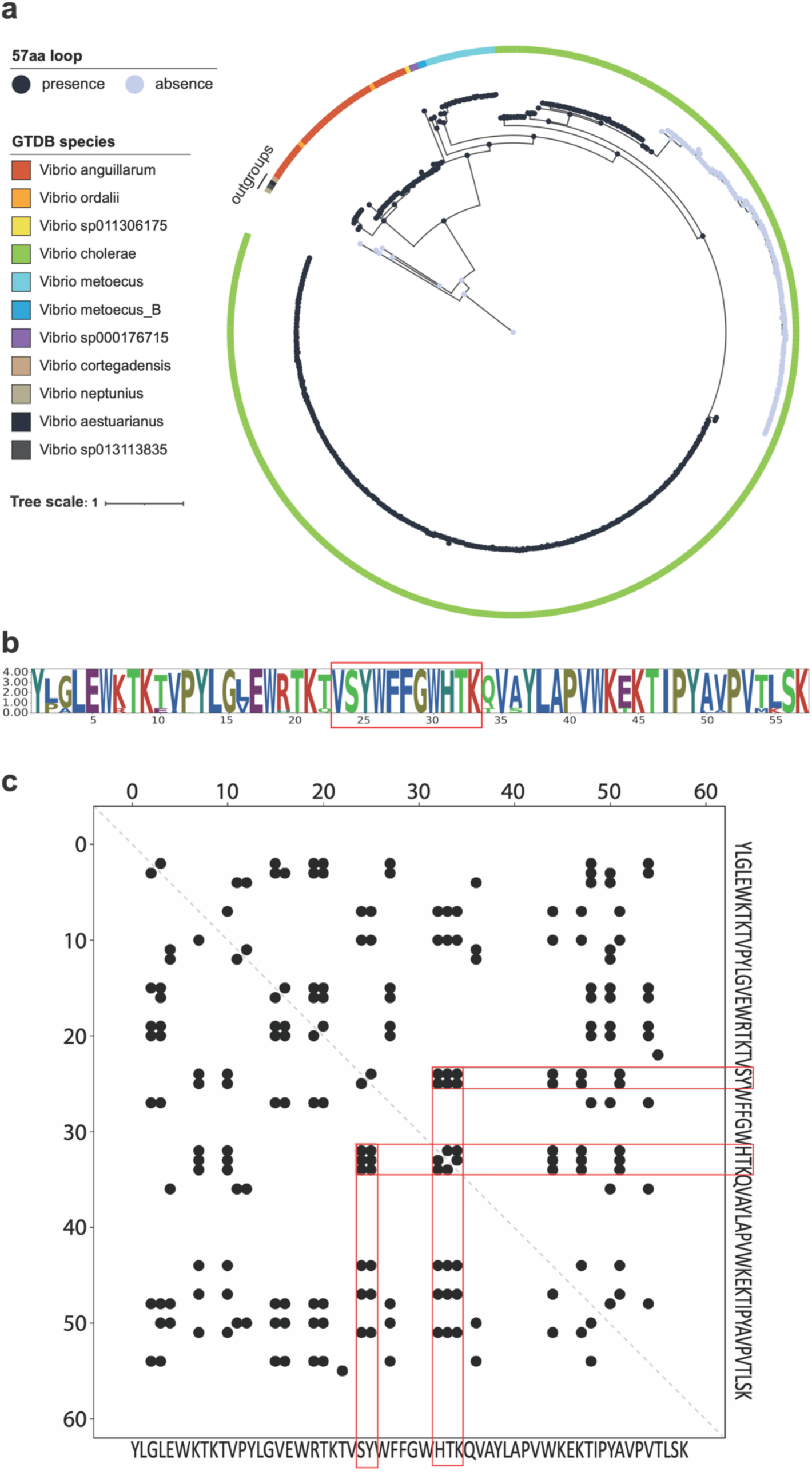
Distribution and conservation of Bap1-57aa across the Vibrio genus. (**a**) Distribution of Bap1-57aa across the Bap1 encoded gene tree. The gene tree is rooted with four RbmC-encoded genes as outgroups. The presence or absence of Bap1-57aa in each gene is denoted by dark blue and light gray circles at the tree tips, respectively. Ancestral states for the presence or absence of Bap1-57aa are shown at internal nodes. A color bar indicates the species origins of the Bap1 encoded genes. (**b**) Sequence logo of Bap1-57aa. The X-axis represents amino acid positions, while the Y-axis represents information content, indicating amino-acid conservation within the sequence. (**c**) Contact map and potential co-evolved residues within Bap1-57aa. The X-axis and Y-axis represent amino acid positions. Black points represent potential co-evolved residues identified using EVcouplings. Red boxes highlight residues two positions upstream and three positions downstream of W_440_FFGW_444_, which are predicted to co-evolve.

Next, we performed a conservation analysis on the 57aa loop among all genes encoding Bap1 with the loop and found that most residues in Bap1-57aa are highly conserved (Fig. 7b). In particular, the core motif, which is suggested to insert into lipid membranes, exhibits extremely high sequence conservation. In contrast, several residues outside the core motif show lower conservation levels. We also conducted a co-evolution analysis using EVcouplings (*52*). Interestingly, this analysis predicts that SY residues at the N-terminus of the core motif co-evolve with HTK residues at the C-terminus of the core motif (Fig. 7c), which may arise from the β-hairpin conformation in which the Y and T form backbone hydrogen bonds (Fig. 4a inset). The strong conservation in the core residues and the predicted co-evolution pattern are consistent with their importance in preserving the structure and functionality of the core motif, as demonstrated by the experimental and simulation results.

## Discussion

In this study, we employed a combination of microscopy, bacterial genetics, and biophysical approaches to unravel the molecular mechanisms of peptide-lipid interactions of a new biofilm-derived peptide. We find that the unique peptide originally discovered in the *Vibrio cholerae* biofilm adhesin Bap1 uses a coordinated strategy for lipid-anchoring: an evolutionarily conserved central segment inserts into lipid membranes in a β-hairpin conformation, while the peripheral sequence, characterized by a pseudo-repeating pattern, facilitates lipid-binding through avidity effects (Fig. 8). Understanding these mechanisms not only advances our knowledge of *V. cholerae* biofilm adhesion but also enriches our general understandings of peptide-lipid binding, particularly in the context of host-microbe interactions. This knowledge may open new avenues for developing bio-inspired adhesives and biofilm control strategies.

**Figure 8.**
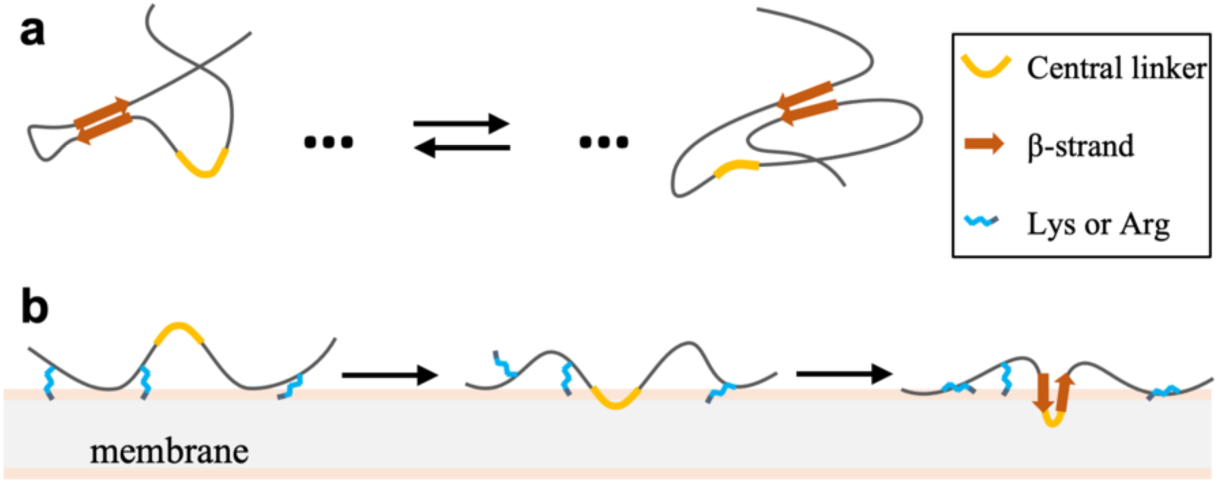
Proposed model for conformational changes and sequence-specific interactions of Bap1-57aa with lipid bilayers. (**a**) Conformational ensemble in solution, featuring transient parallel or antiparallel β-sheets. The chain is in a relatively collapsed state due to intramolecular interactions. (**b**) Pathway for tight membrane binding of Bap1-57aa. After the initial contact through the peripheral repeats, the central linker moves toward the membrane hydrophobic core. This positioning brings the downstream and upstream residues together to form a β-hairpin. β-hairpin formation in turn leads to deeper insertion of the central linker and tighter membrane binding for the entire chain.

Our findings could potentially inspire new ideas for designing lipid-binding motifs with unique structural characteristics. For example, a well-studied common motif in various proteins for membrane binding is the amphipathic helix, which typically relies on bulky hydrophobic residues to detect and bind persistent membrane packing defects, such as those on lipid droplet surfaces (*53*). In contrast to the more common amphipathic helices, β-hairpins have been much less reported to engage in lipid-binding. A recent example is perforin-2 (PFN2), which is thought to enable the killing of invading microbes engulfed by macrophages and other phagocytes (*54*). It was found that the membrane-binding domain in PFN2 employs an extended β-hairpin structure to direct membrane recognition, and that the membrane binding function is primarily conferred by the tip region of the β-hairpin, in a manner conceptually similar to the Bap1-57aa studied here. The conformational changes of the Bap1-57aa peptide show further intricacy in lipid binding—it will be intriguing to ask if similar conformational adaptation also takes place in other membrane-interacting β-hairpins.

Our findings answer several key questions about Bap1-57aa-lipid interaction but also raise new ones. For instance, we observed that the Bap1-57aa peptide tends to form aggregates, yet the underlying mechanism remains unclear. Previous studies have shown that diphenylalanine can self-assemble into various structures, such as tubules and nanowires, depending on the solvent-peptide interactions and the specific sequence (*55–57*). Insertion of residue F_19_ into the hydrophobic core may help stabilize a small Aβ40 oligomer at the membrane surface (*9*). Upon release to solution, the small oligomer can seed fibril growth. While our MD simulations on Bap1-57aa provide insights at the single-chain level, how these peptide molecules interact with each other to form larger structures or complexes, in solution and at a membrane surface, warrants further study.

Another intriguing aspect of lipid-peptide interactions is the role of membrane curvature. Curvature significantly affects the energy landscape of lipid bilayers and their interactions with lipid-binding molecules. Proteins like BAR domains and septin are known to recognize and bind to curved membranes, facilitating processes like cell signaling and division (*58*, *59*). Similarly, bacterial proteins such as MinD use amphipathic helices to detect subtle variations in membrane curvature or composition (*60*, *61*). We have presented evidence showing a preference of the biofilm-derived peptide for sub-40-nm vesicles over larger ones. This suggests a potential curvature-dependent interaction between Bap1-57aa and lipids, which may likely result from higher disorder and more defects of curved membranes. A more refined assay combining lipid composition analysis with precise curvature measurements is necessary to confirm and expand on these observations in the future. This preference could be beneficial for *V. cholerae* cells during colonization: after penetrating the mucosal layer, *V. cholerae* cells may be in direct contact with the intestine epithelial layer, so the ability of the Bap1-57aa to sense the membrane curvature could enable the *V. cholerae* biofilms to preferentially bind the tip of the microvilli that have a diameter ranging from 50-550 nm (*47*).

The dual binding properties on both lipid membranes and abiotic surfaces of the Bap1-57aa peptide open intriguing possibilities for engineering multifunctional adhesives (*21*). Many abiotic surfaces in real-world environments are contaminated with lipid residues or biomolecular films, creating a complex interface. Leveraging the Bap1-57aa peptide’s ability to interact with both lipid and non-lipid substrates could enable the design of multifunctional underwater glues that achieve strong adhesion under diverse challenging conditions. Such adhesives could find widespread applications in biomedical fields, particularly for binding wet surfaces, repairing tissues, or stabilizing implants. Additionally, they could be employed in industrial settings for surface modification, underwater assembly, or biofouling prevention. Further finetuning of the peptide’s sequence or combining it with other functional domains may broaden its utility as a versatile biomolecular tool.

Finally, we put our results into the context of evolution. Bioinformatic analyses reveal that the Bap1-57aa sequence and its surrounding β-prism domain are conserved among several *Vibrio* species (*51*), suggesting a shared role in biofilm adhesion and lipid-binding across these organisms. Interestingly, we found that some isolates have an additional copy of Bap1 without the 57aa sequence, likely resulting from tandem duplication of *bap1*, with the 57aa sequence subsequently lost in one copy. Moreover, the homologous adhesin RbmC has the β-prism domain but not the 57aa loop. These observations suggest that the Bap1-57aa could be considered a genetic element that varies through evolution, being gained or lost as a single unit. Furthermore, the Bap1-57aa sequence itself may have evolved through a series of duplication events to form the four pseudo repeats, as evidenced by the comparison between base pair sequences of the extant sequence and the ancestor sequence (Fig. S7). A directed evolution experiment mimicking the evolutionary pressure imposed on the Bap1-57aa may lead to better lipid-binding peptides for future applications and mechanistic studies.

## Materials and Methods

### Bacterial strains

All *V. cholerae* strains used in this study were derivatives of the wild-type *V. cholerae* O1 biovar El Tor strain C6706str2 and listed in Supplementary Table 1. The rugose strain background harbors a missense mutation in the *vpvC* gene (*vpvC*^W240R^) that elevates intracellular c-di-GMP levels (*62*). The rugose strains form robust biofilms and thus allow us to focus on the biochemical mechanisms governing lipid binding rather than mechanisms involving gene regulation. Additional mutations were genetically engineered into this *V. cholerae* strain using the natural transformation (MuGENT) method (*63*).

### Bacterial growth

All strains were grown overnight in lysogenic broth (LB) at 37°C with shaking. 1× M9 salts were filter sterilized and supplemented with 2 mM MgSO_4_ and 100 µM CaCl_2_ (abbreviated as M9 medium below). Biofilm growth was generally performed in M9 medium supplemented with 0.5% glucose.

### Strain construction

Linear PCR products were constructed using splicing-by-overlap extension (SOE) PCR as previously described and used as transforming DNA (tDNA) in chitin-dependent transformation reactions (*63*). Briefly, SOE PCR was performed by amplifying an upstream region of homology and a downstream region of homology. The desired mutations were incorporated into the primers used in amplification. All primers used to construct and detect mutant alleles are listed in Supplementary Table 2. For chitin-dependent transformation, individual *V. cholerae* colonies were grown in LB media at 30°C for 6 hours to an optical density at 600 nm (OD_600_) = 0.8-1.0. Cells were washed with Instant Ocean (IO) solution and then incubated with chitin particles suspended in IO for 8-16 hours at 30°C before the tDNA was added. The cultures were then incubated at 30°C for an additional 8-16 hours. LB was added to the cultures and incubated at 37°C for 2 hours before plating on LB agar with the appropriate antibiotic. The desired mutants were selected by the emergence of a new phenotype or colony PCR screening and confirmed by sequencing.

### Lipid-coated microbead adhesion assay

Silica microbeads were coated with lipid layers according to published protocols with modification (*29*). Briefly, 100 mol% DOPC (Avanti Polar Lipids 850375), and >0.1 mol% L-α-phosphatidylethanolamine-N-(lissamine rhodamine B sulfonyl) (abbreviated as RhPE, Avanti Polar Lipids 810146) were mixed in chloroform in a glass vial prerinsed with chloroform. A light stream of nitrogen was used to remove excess solvent, followed by at least 2 h in a vacuum desiccator. Lipids were hydrated for 30 min at 37°C at a final lipid concentration of 5 mM in buffer (20 mM Tris, pH 8.0, 300 mM KCl, and 1 mM MgCl_2_) with vortexing and agitation roughly every 5 min and probe sonicated to clarity (4 min, with intermittent breaks) to form small unilamellar vesicles (SUVs). SUVs were adsorbed onto 5 µm silica microspheres by mixing 50 nmol lipids with 440 mm^2^ of silica microspheres surface area in a final volume of 80 µL and 1 h rotary shaking at room temperature. Excess SUVs were removed by pelleting coated beads for 30 s at 862 × g followed by washing 4 times with excess buffer (100 mM KCl and 50 mM Tris, pH 8.0). 100 µL buffer (100 mM KCl, 50 mM Tris, pH 8.0, 0.1% methylcellulose (Sigma-Aldrich M7027), 0.1% BSA) containing FITC-labeled peptide at various concentrations and 0.01 wt% lipid-coated beads was incubated for at least 1 h at room temperature before transferring to a NaOH-treated 96-well plate with a glass bottom. The surface of the 96-well plate was treated with NaOH to render it more hydrophilic and negatively charged. Briefly, before adding the solution, 100 µL of 10M NaOH aqueous solution was added to the wells and incubated at room temperature for 10 min, after which the wells were washed with DI water until the pH was neutral. Thus-prepared samples were imaged with a spinning disk confocal microscope (Nikon Ti2-E connected to Yokogawa W1) using a 60× oil objective (numerical aperture = 1.40) and a 488 nm laser excitation or a 561 nm laser excitation. For each sample, at least three locations were imaged and captured with a sCMOS camera (Photometrics Prime BSI). Each field of view contained roughly 100-150 beads.

### Quantification of bead adsorption assay

The background signal due to the camera in the 488 nm channel was measured by taking images of M9 medium and quantifying them with built-in functions of the Nikon Element software. After subtracting the background signal, the signal intensity per unit area on the surface of the beads and in the solution was calculated using custom MATLAB codes and the difference was determined to give the excess surface signal. The adsorption curve was fitted to a classical Hill binding model.

### CD spectroscopy

Lipid SUVs were prepared as previously described with minor changes. Briefly, 75 mol% DOPC (Avanti Polar Lipids 850375), and 25 mol% DOPS (Avanti Polar Lipids 840035) were mixed in chloroform in a glass vial prerinsed with chloroform. This lipid composition was used due to its optimal optical performance in CD (nearly transparent above 190 nm); we have also verified that the conclusion does not depend on the lipid composition. A light stream of nitrogen was used to remove excess solvent, followed by drying overnight in a vacuum desiccator. Lipids were hydrated for 30 min at 37°C at a final lipid concentration of 5 mM in water with vortexing and agitation roughly every 5 min and probe sonicated to clarity (3-4 min, with intermittent breaks) to form small unilamellar vesicles (SUVs). 100 μM core motif peptide or 50 μM Bap1-57aa peptide was suspended in 4 mM Tris buffer, 10 mM NaCl pH 7.4 and shaking for at least 1 hour. Then the suspension was bath sonicated with ice (6 min, with intermittent breaks) before mixing with different concentrations of lipid SUVs at 1:1 volume ratio. The mixture was incubated for 20 minutes at room temperature before transferring to a CD cuvette. CD wavelength scans were recorded on an Applied Photophysics chirascan circular dichorism spectrometer. All measurements were obtained using a 1-mm pathlength cuvette. Wavelength scans were recorded with an average of 3 repeats, a bandwidth of 2 nm, and a scan rate of 50 nm/min.

### Fluorescence spectroscopy

Fluorescence spectra were recorded on a TCSPC Horiba Fluorolog-QM fluorimeter. All measurements were obtained using a 1.0 cm pathlength cuvette. The sample was excited at 280 nm and monitored from 290 nm to 450 nm.

### Contact angle measurement

*V. cholerae* strains were streaked on LB plates containing 1.5% agar and grown at 37°C overnight. Individual colonies were inoculated into 3 mL of LB liquid medium containing glass beads, and the cultures were grown with shaking at 37°C to mid-exponential phase (5-6 h). Subsequently, the cells in the cultures were vortexed, the OD_600_ was measured, and the cultures were back diluted to an OD_600_ of ∼0.5. 50 µL of this inoculum was applied to an agar plate and spread with a sterile glass rod to enable growth of a biofilm lawn covering the entire plate. Plates were incubated at 37°C for 24 hours to form a continous bacterial lawn. A strip of biofilm (about 1 cm × 4 cm) with the underlying agar was cut out with a razor blade and transferred onto a piece of glass for imaging. To overcome uptake of water by the underlying biofilm/agar, we used a dynamic sessile drop method (*32*): Water was slowly added to the surface by a syringe pump, and the advancing contact angle was measured to approximate the equilibrium contact angle. Side views of biofilm-liquid interfaces were recorded with a Nikon camera (D3300) equipped with a macrolens (Sigma). The contact angle was extracted using the Droplet_Analysis plugin in ImageJ. <u>Biofilm-based adsorption assay:</u> Overnight cultures of the indicated strains constitutively expressing mNeonGreen were grown at 37°C with shaking in 1.5 mL LB. 50 µL from each culture was used to inoculate 1.5 mL of M9 medium supplemented with 0.5% glucose and grown at 30°C with shaking for 5 hours. The inoculant was then bead bashed using a Digital Disrupter Genie with small glass beads (acid-washed, 425 to 500 μm; Sigma-Aldrich). This procedure ensured that large cell clusters formed in culture were broken apart to allow more accurate measurement of OD_600_. The cultures were then diluted to an OD_600_ ≅ 0.5. 100 μL of the regrown culture was aliquoted into a 96-well plate with a glass bottom (MatTek P96G-1.5-5-F) and incubated at 30°C for 1 hour. The wells were then washed twice with M9 medium and replaced with M9 medium with 0.5% glucose. The lid was secured with parafilm and the 96-well plate was subsequently incubated at 30°C for 16 hours. The medium was then replaced by M9 medium with 0.5 mg/mL BSA and 0.1 mM Rhodamine-labeled lipid SUVs. After incubation at room temperature for 1 hour, the wells were washed twice with M9 medium and replaced with M9 medium. Thus-prepared samples were imaged with a spinning disk confocal microscope (Nikon Ti2-E connected to Yokogawa W1) using a 60× water objective (numerical aperture = 1.20) and a 488 nm laser excitation or a 561 nm laser excitation. For each sample, several locations with 2×2 tiles where imaged and captured with a sCMOS camera (Photometrics Prime BSI). The *x*-*y* pixel size was 0.22 μm and the *z*-step size was 1 μm. All images presented in this study are raw data rendered using the Nikon Elements software. For secretion level comparison, the indicated strains tagged with 3×FLAG at the C-terminus and constitutively expressing mNeonGreen were used, and Rhodamine-labeled lipid SUV was substituted by 2 µg/mL anti-FLAG antibody conjugated to Cy3 (Sigma-Aldrich A9594). All other procedures remained the same.

### Quantification of biofilm-based adsorption assay

Image stacks for quantifying SUV-capturing ability or protein secretion level were analyzed using the built-in functions of the Nikon Element software. First, background noise in the 561 nm channel was measured by taking images with M9 medium and subtracting them from the data. Next, image analysis was performed by resizing and thresholding each image layer-by-layer, and measuring the total PerimeterContour above the threshold in each layer. The PerimeterContour for each sample *z*-slice was then summed to give the total biofilm surface area. Subsequently, Rhodamine-labeled SUV signals or anti-FLAG-Cy3 signals were calculated and integrated; the ratio between the total staining signal and the total biofilm surface area was calculated to quantify the ability of a biofilm to capture lipid SUVs (SUV signals) or the protein secretion level (anti-FLAG-Cy3 signals). To diminish interference from the glass substratum, all analyses were performed in the z-range from 1 µm to 36 µm away from the glass substratum.

### *In situ* biofilm immunostaining (SUV colocalization)

Overnight cultures of the indicated strains with WT *bap1* tagged with 3×FLAG at the C-terminus and constitutively expressing SCFP3A were grown from individual colonies at 37°C with shaking in 1.5 mL LB. 50 µL from each culture was used to inoculate 1.5 mL of M9 medium supplemented with 0.5% glucose and grown at 30°C with shaking until the OD_600_ was between 0.1 and 0.3. The cultures were then diluted to an OD_600_ ≅ 0.001. 100 μL of the regrown culture was aliquoted into a 96-well plate with a glass bottom (MatTek P96G-1.5-5-F) and incubated at 30°C for 30 minutes. The wells were washed twice with M9 medium; subsequently, 100 µL of M9 medium with 0.5% glucose, 0.5 mg/ml BSA (Sigma-Aldrich A9647) and 2 µg/mL anti-FLAG antibody conjugated to FITC (Sigma-Aldrich A9594) was added to the well. The lid was secured with parafilm and the samples were incubated at 30°C for 40-48 hours. The medium was then replaced with M9 medium with 0.5 mg/mL BSA and 0.1 mM previously prepared Rhodamine-labeled lipid SUV. After incubation at room temperature for 1 hour, the wells were then washed twice with M9 medium and replaced with M9 medium. Thus-prepared samples were imaged with a spinning disk confocal microscope (Nikon Ti2-E connected to Yokogawa W1) using a 100× oil immersion objective (numerical aperture = 1.35), a 445 nm laser excitation to observe the cells, a 488 nm laser excitation to observe protein localization, and a 561 nm laser excitation with the corresponding filters to observe lipid SUVs. The images were captured with an sCMOS camera (Photometrics Prime BSI) at a *z*-step size of 0.5 µm.

### Caco-2 cell culturing and staining

Human colonic epithelial Caco-2 cells (ATCC HTB-37) were obtained from ATCC and authenticated by ATCC based on morphology, doubling time, and STR profiling. Caco-2 cells were cultured in flasks containing Dulbecco’s Modified Eagle’s Medium (DMEM; Gibco) supplemented with 10% (v/v) heat-inactivated fetal bovine serum (FBS-HI; Gibco) at 37 °C in a humidified 5% CO_2_ incubator. After 72 h, cells were collected via dissociation using 0.25% Trypsin-EDTA (Gibco) and pelleted by centrifugation (300 rcf, 3 min, room temperature in 15 mL conical tubes (Corning); then 21,000 rcf, 2 min, room temperature in Eppendorf tubes). Cell pellets were stored at −80 °C prior to further analysis.

To stain Caco-2 cells with purified proteins, a frozen aliquot of Caco-2 cells as prepared above was gently thawed and then added to 1.5 mL of M9 medium containing 1 mg/mL BSA, 300 nM DAPI and 0.11 µM Alexa Fluor 647 phalloidin (Invitrogen, A22287) and incubated for 15 min at room temperature. 100 µL of this cell suspension was aliquoted to sterile 1.5 mL microcentrifuge tubes and spun at 500 × g for 5 min. The staining media were removed and replaced with 100 µL of M9 media containing 1 mg/mL BSA and 1.5 µM of FITC-labeled peptides. The samples were incubated for 30 min at room temperature and then the media was replaced with 100 µL fresh M9 medium and transferred to the wells of a 96-well plate. The samples were imaged with a spinning disk confocal microscope using a 60× oil objective and a 405 nm laser excitation to observe the Caco-2 cell nuclei and a 488 nm laser excitation to observe peptide localization, a 561 nm laser excitation to observe membrane or a 640 nm laser excitation to observe actin skeleton with the corresponding filters. Image stacks were taken from 2 μm to 30 μm to reduce signal interference from substrate layer.

### *bap1* gene identification

Among the 1,983 genomes we identified from the Vibrio genus in the GTDB r214 (Genome Taxonomy Database) (*64*), we selected only those with completeness ≥ 90% and contamination ≤ 5% for further analysis. To eliminate genome redundancy, we employed Mash v2.3 (*65*) to determine pairwise distances between genomes (using a threshold of -d 0.01) and grouped them using Markov Clustering (*66*), resulting in 413 distinct clusters. A representative genome was selected from each cluster. We identified 4,066 RbmC and Bap1 encoded sequences by querying WP_000200580.1 (RbmC) and WP_001881639.1 (Bap1) against all protein sequences in the genomes using BLASTp v2.15.0+ (*67*), with criteria of > 40% identity, > 250 bit score, and > 200 amino acids in aligned length.

### *bap1* encoded gene tree construction

We performed multiple sequence alignments using high-quality RbmC and Bap1 genes, defined as those with ≥80% identity to a Bap1 query, lengths of 650–700 aa, and bit scores >900, after removing sequence redundancy. We applied MAFFT v7.475 (*68*) to align high-quality protein sequences with options “--maxiterate 1000 --localpair” and aligned low-quality protein sequences by adding them to the previously aligned high-quality genes using MAFFT with option “-add”. The aligned protein sequences were mapped back to the nucleotide sequences to align by codons using PAL2NAL v14 (*69*). Finally, a codon-based phylogenetic Bap1 encoded gene tree containing 392 sequences was built with the aligned nucleotide sequences using RAxML v8.2.12 (*70*) by providing a partition file (“-m GTRGAMMA -q dna12_3.partition.txt”).

### Bap1-57aa loop extraction and ancestral sequence inference

The domain boundaries of all Bap1 and RbmC encoded genes were manually determined by investigating the multiple sequence alignment in Geneious Prime v2023.1.2 (https://www.geneious.com) and cross-validated using predicted structures generated by ESMfold v2.0.0 (*71*). After domain segmentation, the Bap1-57aa loops were extracted from Bap1-encoded genes. The presence or absence of a Bap1-57aa loop was then mapped to the terminal nodes of the Bap1 encoded gene tree, and their ancestral sequences, along with the inferred presence or absence states, were reconstructed using asr.GRASP (*72*).

### Vesicle preparation for flotation assays

To make SUVs and LVs, a lipid mixture containing 84.2% DOPC, 15% DOPS and 0.8% Cy5-DOPE (Avanti Polar Lipids) was first dried by nitrogen gas. Any remaining organic solvent was removed by placing the lipid film under vacuum overnight. The lipid film was hydrated with water to a 5 mM stock and shaken using an Eppendorf Thermomix for >30 min and stored at −20 °C. Before usage, the lipid suspension was diluted to 200-300 μM in Buffer A (10 mM HEPES pH 7.4, 150 mM NaCl). After shaking for >30 min, the lipid suspension was then split into two tubes. To prepare LVs, the lipid suspension was extruded 19 times through a 400-nm pore size polycarbonate filter (Whatman) using a mini-extruder (AVESTIN). For SUV preparation, the lipid suspension was sonicated on ice for 2 min at 20% amplitude, with 2 s sonication followed by 4 s pause, using a Q125 Sonicator with a 1/8” probe (Qsonica). The SUVs were centrifuged at 20,000 g for 5–10 min to remove metal particles.

### Vesicle-peptide co-floatation assay

Samples (150 μL each) containing 1 μM Bap1-57aa WT peptide and liposomes containing 50 μM lipids were mixed with 200 μL 60% sucrose (m/v% in Buffer A), reaching a final sucrose concentration of ∼34%. The sucrose-containing samples were then layered at the bottom of 5 mm × 41 mm Beckman ultracentrifuge tubes (#344090) and overlaid with 200 μl of 25% sucrose (in Buffer A) and finally 100 μl of Buffer A (Fig. 6). The tubes were spun for 1 hour at 48k rpm and 4 °C in a SW 55 Ti rotor. Fractions were collected as shown in Fig. 6, and their contents were determined by SDS-PAGE using two-layer (4% over 10%) stacking gels made in house.

### Negative stain TEM study and vesicle size measurement

A drop of sample from flotation fractions (8 μL) was deposited on a glow discharged formvar/carbon-coated copper grid (Electron Microscopy Sciences), incubated for 2–3 minutes and blotted away. The grid was then washed briefly and stained for 1 min with ∼ 5 μL of 2% (w/v) uranyl formate. Images were acquired on a JEOL JEM-1400 Plus microscope (acceleration voltage: 80 kV) with a bottom-mount 4k×3k CCD camera (Advanced Microscopy Technologies) using the AMT Image Capture Engine. Vesicle sizes were measured from electron micrographs by ImageJ (National Institutes of Health) automatically for SUVs (*73*) or manually for LVs. The diameter of each liposome was calculated based on the measured area (A) following the equation: 𝐷 = 2 × &𝐴/π.

### Molecular dynamics simulations of disordered Bap1-57aa in a lipid environment

Disordered conformations of Bap1-57aa were generated using the TraDES method (*25*). Two extended conformations were chosen and placed near a membrane with a composition of PC:PS:PIP_2_ at 75:20:5, same as in our previous study of the membrane association of a protein with a disordered region (*24*) and mimicking the plasma membrane. The CHARMM-GUI web server (*74*) was used to prepare the peptide-lipid systems, with 250 lipids in each leaflet. The force field for the peptide and membrane was CHARMM36m (*75*). TIP3P water (*76*) was used to solvate each system in a cubic box with a 130 Å side length. Na^+^ and Cl^−^ ions were added to neutralize the system and generate a 150 mM NaCl concentration. The total number of atoms was 209,846 and 218,700 with the two Bap1-57aa initial conformations.

Each system was equilibrated using NAMD 3.0 (*77*) following a six-step protocol from CHARMM-GUI. Specifically, after 10,000 steps of conjugate-gradient energy minimization, the first two steps of the equilibration were at constant temperature and volume (NVT) (each for 125 ps), and the last four steps were at constant temperature and pressure (NPT) (for 125, 500, 500, and 500 ps). Constraints on lipid head group and protein backbone were gradually reduced. The timesteps were 1 fs for the first three steps and 2 fs for the remaining steps. Finally, production runs were performed for 1.1 μs in four replicates for each system at constant NPT with a 2 fs timestep using *pmemd.cuda* (*78*) in AMBER 22 (*79*). In three of the eight replicate simulations, Bap1-57aa inserted into the lipid bilayer via the central linker, within the first 100-350 ns. Snapshots after the initial insertion were saved at 100 ps intervals for further analysis.

All bonds connected to hydrogens were constrained by the SHAKE algorithm (*80*). Long-range electrostatic interactions were treated by the particle mesh Ewald method (*81*). The cutoff distance for nonbonded interaction was 12 Å, with force switching at 10 Å for van der Waals interactions. The Langevin thermostat with a damping constant of 1 ps^-1^ was used to maintain constant temperature at 310 K. The Berendsen barostat (*82*) was used to maintain pressure at 1 atm.

A second set of 12 replicate simulations (IDP-wr) was performed, with the Cα-Cα distance between the N- and C-terminal residues restrained to 28 Å. The latter was the mean value in the 8 IDP-wor simulations during the 50-450 ns period. The restraint was harmonic with a force constant of 250 kcal mol^-1^Å^-2^, imposed through PLUMED (*83*). The central linker was initially placed at a similar location to the counterpart during the initial insertion in the IDP-wor simulations. The IDP-wr simulations were also run for 1.1 μs. Snapshots in the last 1 μs were saved at 100 ps intervals for analysis.

### Molecular dynamics simulations of AF-melt Bap1-57aa in the lipid environment

The AlphaFold predicted structure of Bap1 was downloaded from Uniprot (entry A0A7Z7YFH0). The portion for the Bap1-57aa loop (residues 415-471) was taken and placed in a cubic box with a side length of 120 Å. Solvation by TIP3P water and neutralization by Cl^−^ resulted in a total of 165,929 atoms. After a 5000-step minimization using *sander*, a 250-ps equilibration at constant NVT was performed in *pmemd.cuda* at a 1 fs timestep, with positional restraints on the peptide with a force constant of 1 kcal mol^-1^ Å^-2^. The temperature was ramped from 0 to 500 K in the first 50 ps and maintained at 500 K for the remaining 200 ps. A production run was performed at constant NVT (*T* = 500 K) for 100 ns at a 2 fs timestep without restraint. After visual inspection, the snapshot at 11 ns, with an antiparallel β-sheet (two strands, each with four residues), was chosen to prepare AF-melt simulations at the membrane surface. Specifically, this AL-melt structure was placed at a similar location to the IDP-wor counterpart during the initial insertion. The simulation box dimensions were 130 Å ξ 130 Å ξ 155 Å, with a total number of 258,693 atoms. The Cα-Cα distance between the N- and C-terminal residues was restrained to 20 Å with a force constant of 250 kcal mol^-1^ Å^-2^, to help maintain some residual structure. Four replicate simulations were run for 400 ns each. Analysis was done on the last 350 ns.

### Molecular dynamics simulations of Bap1-57aa in a β-hairpin conformation at the membrane surface

Residues S_438_YWFFGWHTK_447_ were modeled as a β-hairpin (Fig. 4a inset) using XPLOR-NIH (*84*) with simulated annealing. F_442_G_443_ were assigned to the *i* + 1 and *i* +2 positions; hydrogen bonds were formed between F_441_ and W_444_ and between Y_439_ and T_446_. Phi and psi angles for β-sheet residues S_438_ to F_441_ and W_444_ to K_447_ were assigned to −140° and 130°, respectively. Simulated annealing was performed from 3500 K to 25 K in 1000 steps, and the force constants for the restraints were ramped up from 5 to 1000 kcal/mol rad^-2^ for phi and psi angles and from 2 to 30 kcal/mol Å^-2^ for backbone hydrogen-bond donor-acceptor distances. The rest of Bap1-57aa was modeled as disordered using TraDES (*25*), and joined with the β-hairpin using VMD (*85*). The tip of the β-hairpin was initially placed at 4 Å below the phosphate plane, while the disordered regions was away from the membrane surface. The simulation box dimensions were 130 Å ξ 130 Å ξ 125 Å, with 205,687 atoms. Eight replicate simulations were run for 1.1 μs. Analysis was done on the last 1 μs.

### Molecular dynamics simulations of Bap1-57aa in a β-hairpin conformation in solution

The initial structure of Bap1-57aa modeled above was placed in a cubic box with a side length of 119 Å. Solvation by TIP3P water and 150 mM NaCl resulted in a total of 158,927 atoms. A 10,000-step minimization was followed first by a 1-ns equilibration at constant NVT and a 1 fs timestep, with positional restraints on the peptide backbone with a force constant of 5 kcal mol^-1^ Å^-2^, and then by a 2-ns equilibration at constant NPT and a 2 fs timestep without restraint. Finally, four replicate simulations were run for 400 ns at constant NPT. Analysis was done on the last 350 ns.

### Analysis of simulation data

CPPTRAJ (*86*) and in-house python scripts were used to calculate contact frequency, Z_tip_ distance, secondary structure, and end-to-end distance from the trajectories. Lipid contact was calculated with a 3.5 Å cutoff between heavy atoms of a peptide residue and any lipid molecule. Z_tip_ distance was defined as the mean Z coordinate of the tip heavy atom of each side chain, in a Cartesian coordinate system with the Z axis along the membrane normal and X-Y plane at the mean Z coordinate of the phosphorus atoms in the proximal leaflet. After averaging over saved snapshots in each simulation and then over replicate simulations, data were presented as plots. Pymol (https://pymol.org) and VMD were used to render images and to make a movie. <u>Statistics and reproducibility:</u> Error bars correspond to standard deviations from measurements taken from distinct samples. Standard *t*-tests were used to compare treatment groups and are indicated in each figure caption. Tests were always two-tailed, unpaired, and with Welch’s correction, as demanded by the details of the experimental design. All statistical analyses were performed using GraphPad Prism software. Microscopy images and spectra were shown from representative results from at least three independent experiments.

## Data availability

Source data are provided with this paper.

## Materials availability

All bacterial strains constructed as part of this work will be provided to the community upon request in a timely fashion and shipped in accordance with biosafety standards and regulations.

## Acknowledgements

We thank Dr. Longfei Liu for help with liposome preparation and discussions. We thank Ms. Katherine Matej and Dr. Hualiang Pi for providing the Caco-2 cells. We thank Drs. Shoken Lee, Shirin Bahmanyar, Xiaolei Su, and Erdem Karatekin for helpful discussions. We thank Drs. Chih-Hao Lu, Bianxiao Cui, Rana Ashkar, and Suryabrahmam Buti for experimental help on the curvature dependence of lipid-binding. We thank Drs. Merrill Asp and Kee-Myoung Nam for help with MATLAB codes.

## Author Contributions

Conceptualization: XH, HXZ, JY

Strain construction and validation: XH

Bead-based assays performance and quantification: SS, XH

Peptide characterization, biofilm imaging and Caco-2 cell staining: XH

Biophysical characterizations: XH, SOS, CMD

Measurements on curvature sensitivity: QY, CL

MD simulations and MD data analysis: RP

Phylogenetic analysis: YY, XJ

Writing – original draft: XH, RP, YY, QY, CL, HXZ, JY

Writing – review & editing: XH, RP, YY, SOS, RO, CMD, XJ, HXZ, JY

## Data and materials availability

All final data are available in the main text or the supplementary materials. Correspondence and requests for raw data should be addressed to either.

