## Supplementary material for "Conformations and sequence determinants in the lipid binding of an adhesive peptide derived from *Vibrio cholerae* biofilm": SI figures and Tables

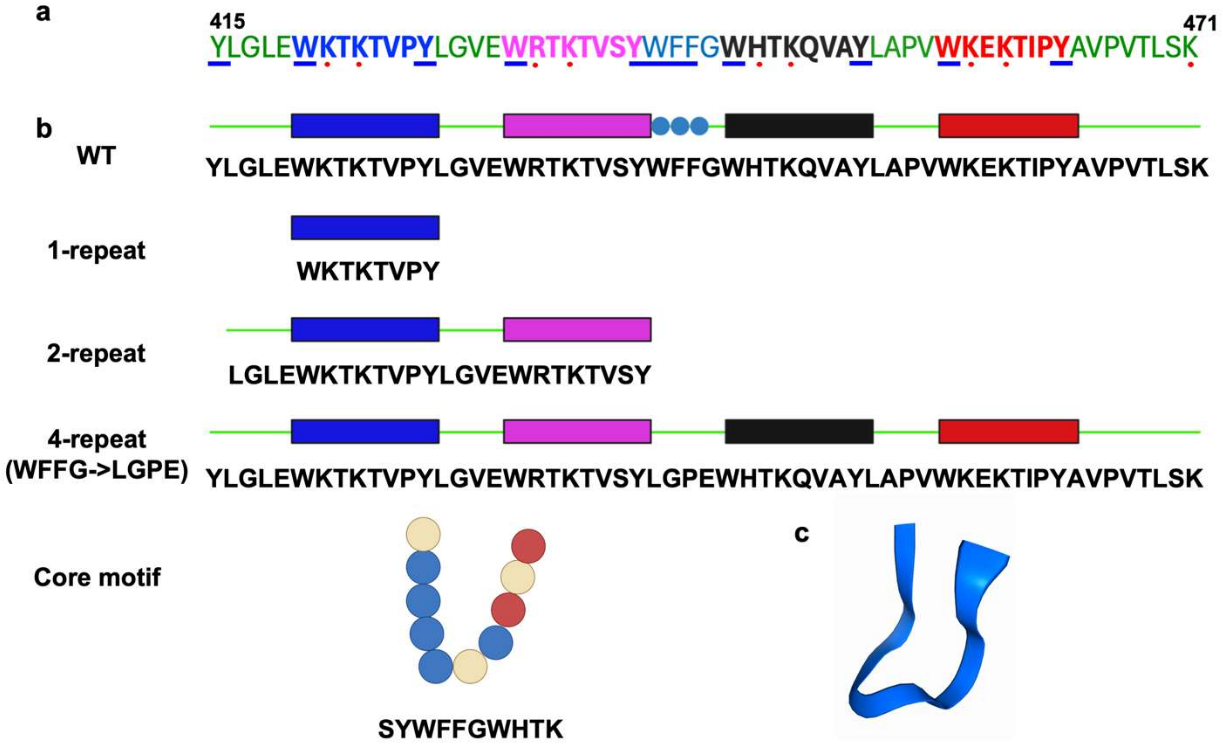


**Fig. S1.**

Schematics and sequences of different peptides used in this study. (**a**) Sequence of WT Bap1-57aa. Aromatic and basic residues are indicated by blue underline and red dots, respectively. Four pseudo repeats are shown in blue, magenta, black, and red. (**b**) Schematic representation and sequences of various peptides in this study. Colored circles indicate amino-acid types: blue, aromatic; red, positively charged; beige, other. (**c**) AlphaFold3 prediction of the core motif peptide (predicted with alphafoldserver.com).

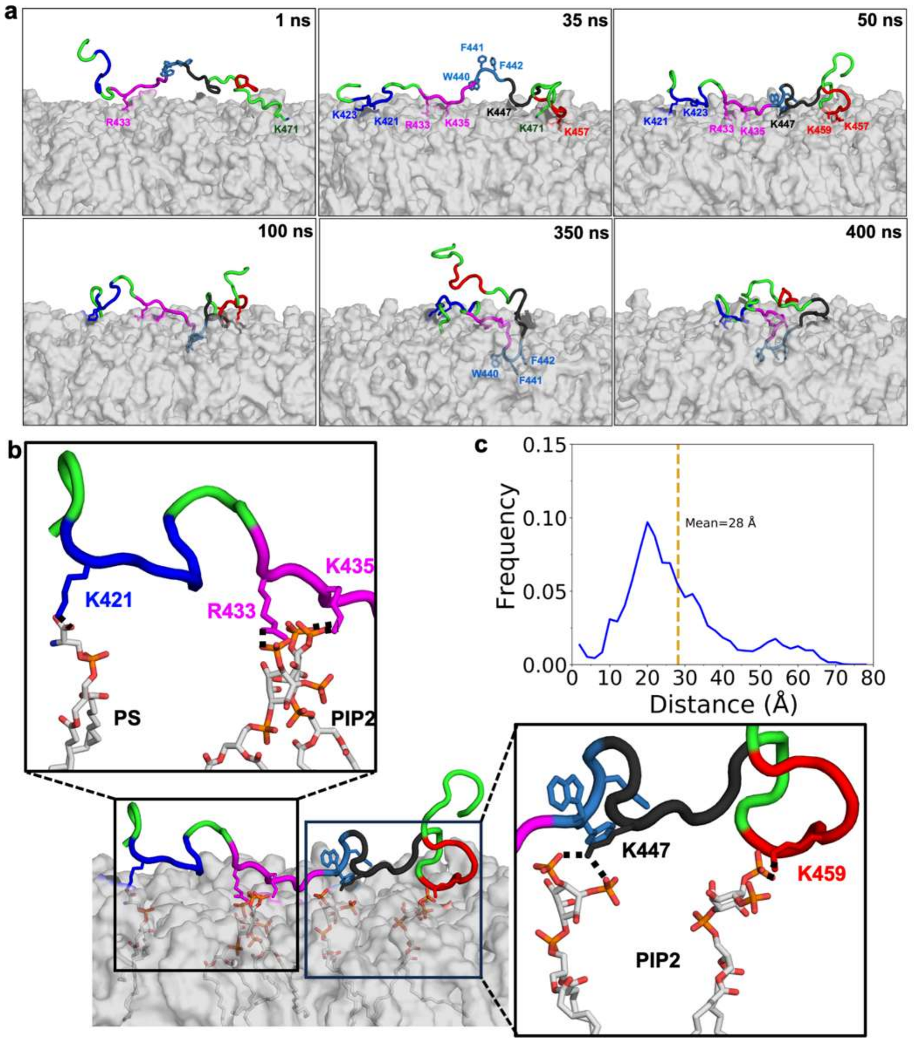


**Fig. S2.**

Simulation results of Bap1-57aa interacting with the membrane in a disordered confirmation without an end-to-end constraint (IDP-wor). (**a**) Snapshots before (1, 35, 50 ns), during (100 ns), and after (350 and 400 ns) the initial insertion of the middle linker WFFG. (**b**) Enlarged view of the snapshot at 50 ns. Multiple basic residues form salt bridges with the headgroups of acidic lipids. (**c**) Distribution of the end-to-end distance from 8 IDP-wor simulations of Bap1-57aa, calculated in the period from 50 to 450 ns.

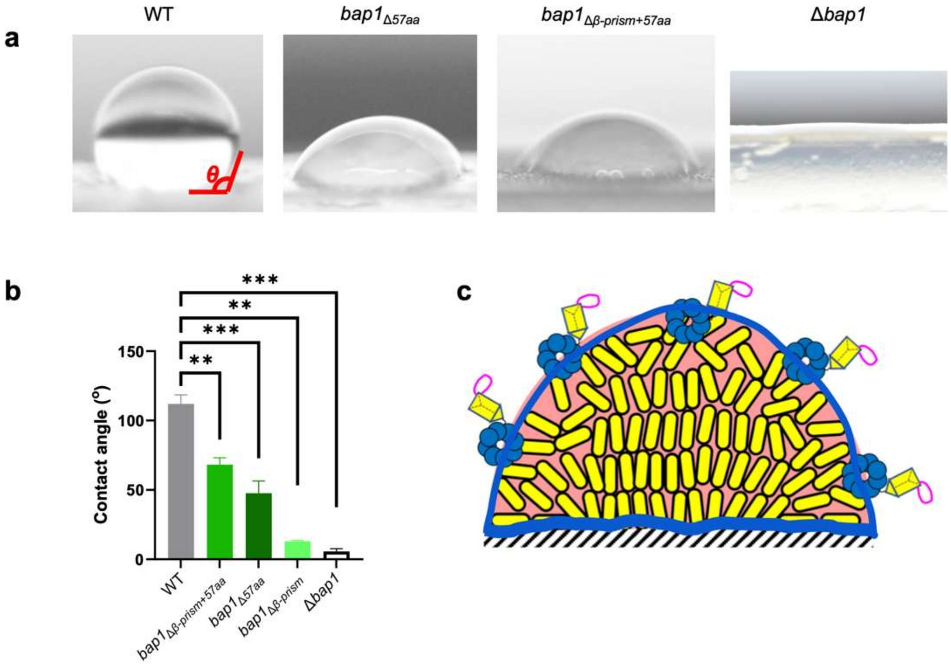


**Fig. S3.**

Contact angle measurements suggest that Bap1 adopts a geometry with the 57aa loop facing outward. (**a**) Image of a water droplet on a biofilm of the indicated strain grown on 1.5% LB agar. All strains are in a Δ*rbmC* background. (**b**) Quantification of the contact angle of water droplets on biofilms of the indicated strains. *V. cholerae* biofilms are hydrophobic, as confirmed by the contact water contact angle measurements. Previous results have shown that this hydrophobicity arises from Bap1(*1*, *2*), specifically the extensive aromatic residues on the cap of β-prism and in the 57aa loop. Data are shown as the mean ± SD of 3 biological replicates. ** $p < 0.01$; *** $p < 0.001$. Exact $p$ values from left to right: 0.0011, 0.0008, 0.0013, 0.0005. (**c**) Schematic suggesting the geometry of Bap1 in biofilms. We hypothesize that the 57aa faces outward to generate the hydrophobicity of *V. cholerae* biofilms. In this geometry, Bap1 molecules are ready to interact with exogenously added lipid SUVs in the assay in Figure 2.

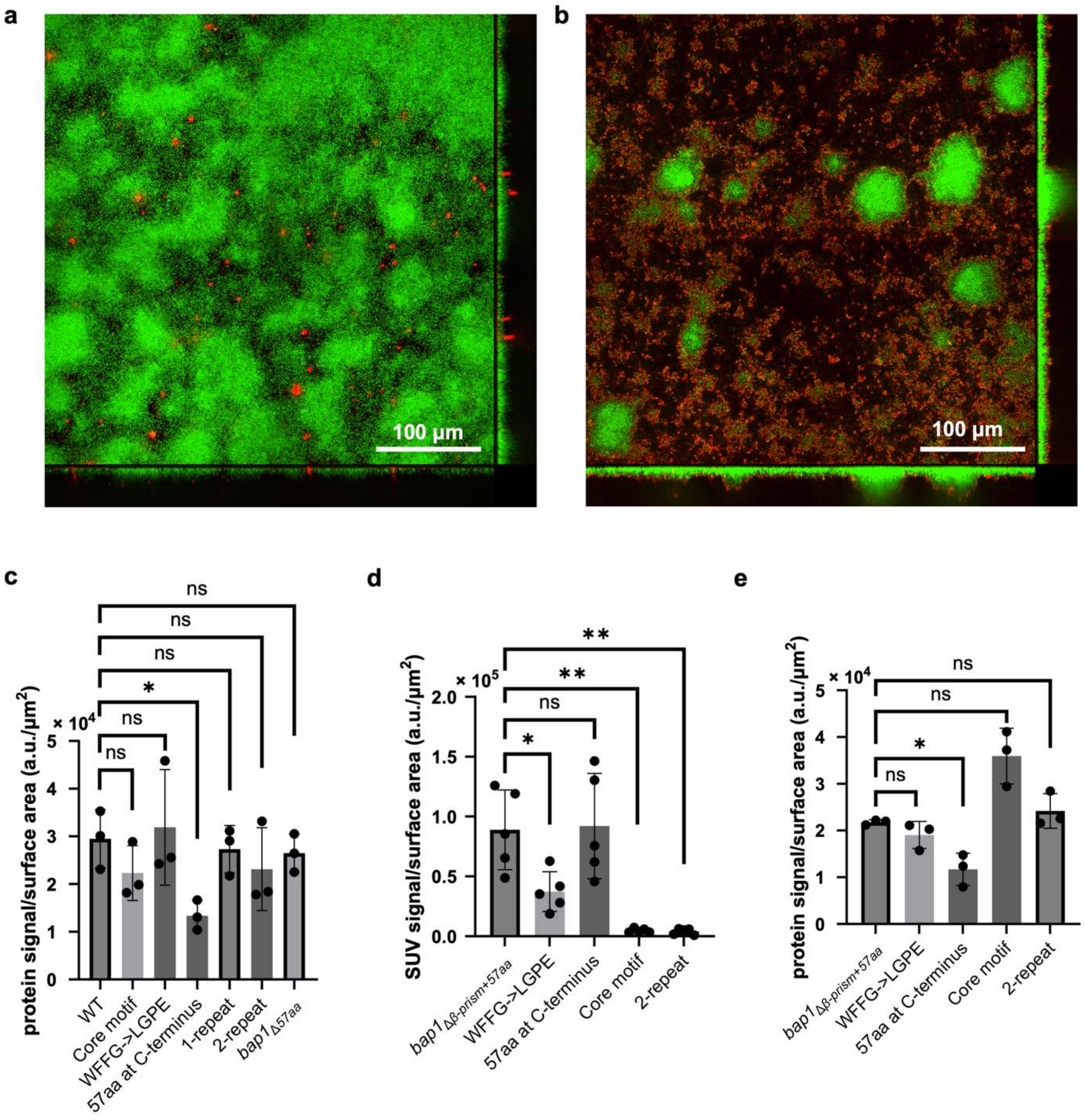


**Fig. S4.**

Bap1 variants show comparable secretion levels. (**a**) Representative image of SUVs (red) adhering to a biofilm grown from cells constitutively expressing mNeonGreen (green) and possessing the WT Bap1-57aa sequence, after a vigorous washing step. (**b**) Representative cross-sectional images (at $z = 9$ μm) of biofilms from cells constitutively expressing mNeonGreen (green) and possessing WT Bap1 tagged with a 3×FLAG epitope at the C-terminus and stained with an anti-FLAG antibody conjugated to Cy3 (red). The secretion levels of Bap1 mutants were quantified by taking the ratio of total signals from an anti-FLAG antibody over the biofilm surface area. (**c**) Quantification of the secretion levels of Bap1 based on assays shown in (b). All data are depicted as the mean ± SD ($n$ = 3 biological replicates). Statistical analysis was performed using two-tailed $t$-test with Welch's correction. ns stands for not significant; * $p < 0.05$. Exact $p$ values from left to right: 0.2123, 0.7804, 0.0271, 0.6570, 0.3628, 0.5220. (**d**) Quantification of SUV capture by biofilms grown from Bap1 mutants in a background in which the β-prism was deleted. The cap of the β-prism contains a few exposed tryptophan and lysine residues, which may also contribute to lipid binding. In the $bap1_{\Delta\beta\text{-}prism}$ background, the WFFG→LGPE mutant showed significantly reduced SUV capture compared to the intact Bap1-57aa. Moreover, the mutant containing only the core motif has abolished the SUV capturing ability. We reasoned that in this strain background, the large β-propeller domain poses a steric hindrance as to prevent the shorter core motif from binding to SUVs. All data are depicted as the mean ± SD ($n$ = 5 biological replicates). Statistical analysis was performed using two-tailed $t$-test with Welch's correction. ns stands for not significant; * $p < 0.05$; ** $p < 0.01$. Exact $p$ values from left to right: 0.0219, 0.5737, 0.9006, 0.0048, 0.0046. (**e**) Quantification of the secretion level for *bap1* mutants in the background with the β-prism removed. All data are depicted as the mean ± SD ($n$ = 3 biological replicates). Statistical analysis was performed using two-tailed $t$-test with Welch's correction. ns stands for not significant; * $p < 0.05$. Exact $p$ values from left to right: 0.2380, 0.1094, 0.0347, 0.0536, 0.3797. a.u. stands for arbitrary unit. For strains tested in (**c**) and (**e**), most Bap1 mutants are secreted at similar levels to WT Bap1 except for the mutant in which the 57aa was moved to the C-terminus of Bap1 in the respective background. Nevertheless, these mutants are fully functional in lipid binding regardless of the strain background.

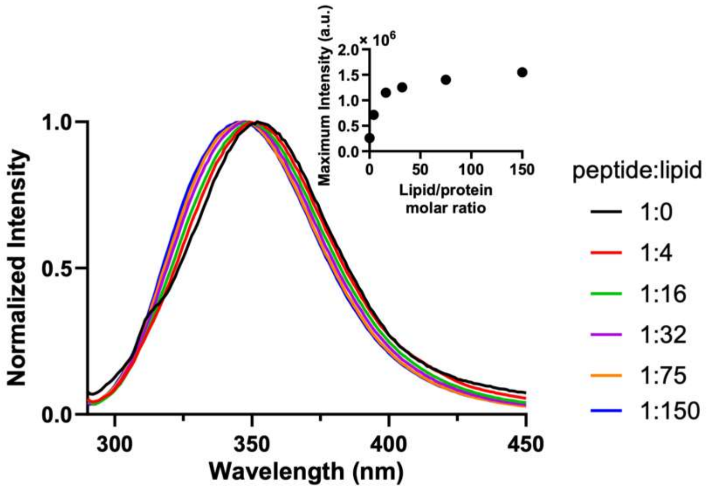


**Fig. S5.**

Normalized tryptophan fluorescence spectra of the WFFG→LGPE peptide mixed with 3:1 DOPC/DOPS SUV at molar ratios between 1:0 and 1:150 in 10 mM Tris buffer pH 7.4 and 150 mM NaCl. Inset: maximum tryptophan fluorescence intensity of the peptide plotted over a series of peptide:lipid molar ratios. Compared with the WT peptide, a similar increase is observed in the emission intensity of the tryptophan residues in the WFFG->LGPE mutant upon lipid binding, but the blueshift of the peak is reduced. The lowest peak position upon lipid binding is 347 nm as compared to 338 nm for the core motif and WT Bap1-57aa. This reduced blue shift suggests that the peripheral tryptophans may not be buried as deeply as the tryptophan residues in the core motif.

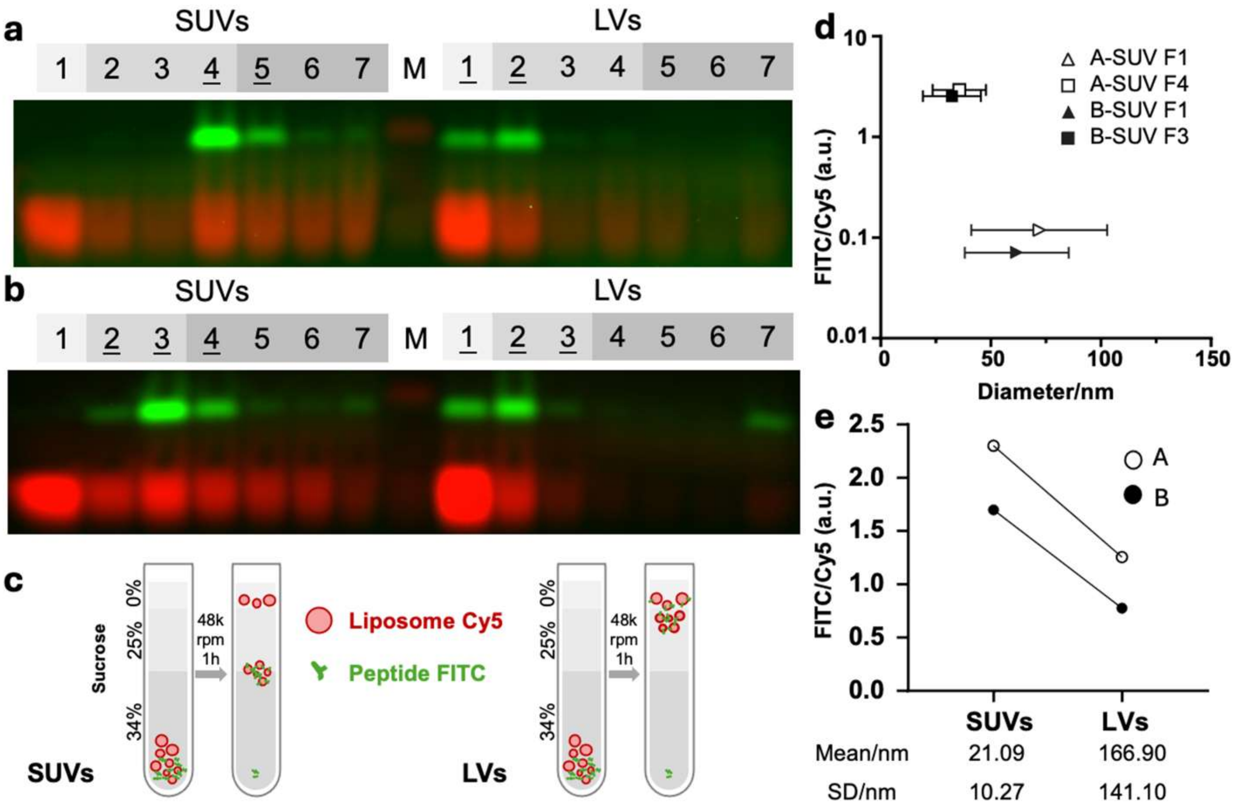


**Fig. S6.**

Bap1-57aa binding to SUVs and LVs analyzed by flotation assays. (**a**) Schematic of the floatation assay. Peptide-coated SUVs and LVs have different densities and thus float to different layers in the gradient. (**b**-**c**) SDS-PAGE analysis of flotation assays with SUVs (prepared by sonication) and LVs (extruded by 400 nm filter). In the two trials, seven fractions were collected from slightly different sucrose gradients. Sucrose concentration and volume of each layer are denoted by gray scale and width of the rectangles, respectively. Pseudo-colors: Cy5, red; FITC, green. (**d**) FITC to Cy5 fluorescence ratio [measured from the gel images in (b) and (c) and normalized to $\sum_{F1}^{F7}(\text{FITC}) / \sum_{F1}^{F7}(\text{Cy5})$ for each experiment] plotted as a function of mean diameter of SUVs (measured by negative-stain TEM) for selected fractions recovered from gradients loaded with SUVs. For each experiment, two fractions with the highest Cy5 fluorescence (i.e., vesicles) are quantified. Error bars show standard deviation, $n = 41$ (F1, trial A), 38 (F4, trial A), 73 (F1, trial B) and 89 (F3, trial B). The SUV part of the gel image in (c) is identical to that in Fig. 6b and shown here for comparison. (**e**) Comparing FITC to Cy5 fluorescence ratios on LVs and SUVs. For each gradient, the FITC and Cy5 fluorescence in fractions containing both peptides and vesicles are quantified; Σ(FITC)/Σ(Cy5) of these fractions [underlined in (b-c)] are normalized to that of all fractions and plotted. a.u. stands for arbitrary unit. Vesicle sizes measured from negative-stain TEM for replicate B before peptide binding are listed below the graph for reference. $n = 474$ (SUV) and 164 (LV).

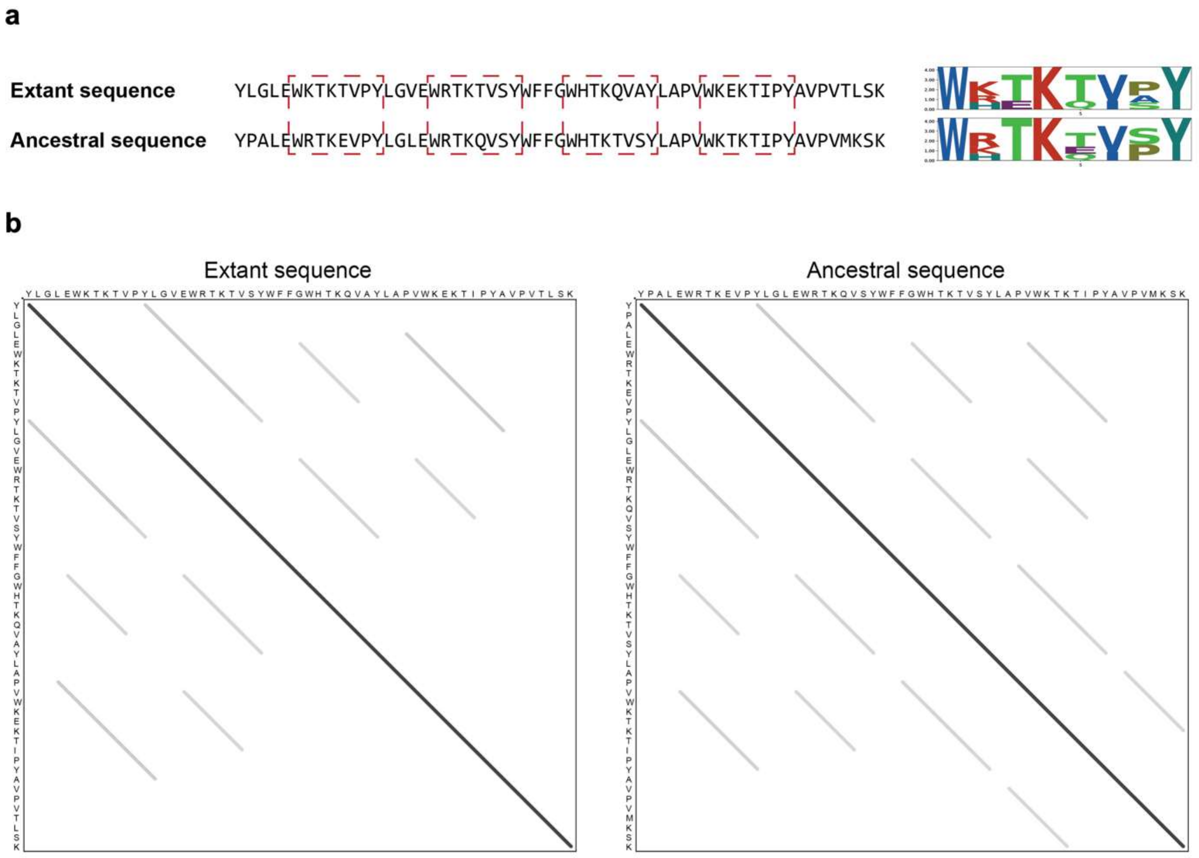


**Fig. S7.**

Comparison between extant and ancestral Bap1-57aa loop sequences. (**a**) Comparison of the repeat regions (highlighted in red boxes) between the extant and ancestral 57-aa loops. The extant sequence is derived from the *V. cholerae* O1 biovar El Tor str. N16961 (accession: GCA_000006745.1). The ancestral sequence was reconstructed using asr.GRASP at the outermost internal node where the 57-aa loop is predicted to have first appeared during the evolution of the Bap1 encoded gene (refer to the Bap1 encoded gene tree in Fig. 7a). (**b**) DNA dot plots of the extant and ancestral 57-aa loops. Lines parallel to the diagonal indicate the repeat regions in the sequences. While repeat patterns are evident in both the extant and ancestral 57-aa loops, the ancestral sequence exhibits enhanced similarity among the repeat regions. This suggests that the Bap1-57aa may have evolved from repeated duplication of a shorter ancestral sequence.

| **Strain Name in Manuscript** | **Genotype and Antibiotic Resistance** | **Description** | **Strain# & Reference** |
|---|---|---|---|
| Rg background | *vpvC*$^{W240R}$, Sm$^R$ | Missense mutation in the *Vibrio cholerae* O1 El tor strain that elevates the level of cyclic-di-GMP. Rugose phenotype. Serves as the parental strain for most of the mutants in this manuscript | JY028 (*3*) |
| Δ*rbmC* | *vpvC*$^{W240R}$, Δ*rbmC*, Δ*VC1807*::*P*$_{tac}$-*mNeonGreen*, Spec$^R$ | Clean deletion of *rbmC* by cotransformation with Δ*VC1807*::*P*$_{tac}$-*mNeonGreen,* Spec$^R$ | ZJ033 (*4*) |
| Δ*rbmC* Δ*bap1* | *vpvC*$^{W240R}$, Δ*bap1,* Δ*rbmC,* Δ*VC1807*::*P*$_{tac}$-*mNeonGreen*, Spec$^R$ | Clean deletion of *rbmC* by cotransformation with Δ*VC1807*::*P*$_{tac}$-*mNeonGreen,* Spec$^R$ into a rugose strain lacking *bap1* (JY074) | STN0009 (*5*) |
| Δ*rbmC bap1*$_{Δ57aa}$ | *vpvC*$^{W240R}$, Δ*rbmC, bap1*$_{Δ57aa}$, Δ*VC1807*::*P*$_{tac}$-*mNeonGreen,* Spec$^R$ | Replacement of *bap1*$^{WT}$ with a *bap1* construct lacking the 57aa by cotransformation with Δ*VC1807*::*P*$_{tac}$-*mNeonGreen,* Spec$^R$ into a rugose strain lacking *rbmC* (JY071) | ZJ032 (*4*) |
| Δ*rbmC bap1*$_{Δβ\text{-}prism+57aa}$ | *vpvC*$^{W240R}$, Δ*rbmC, bap1*$_{Δβ\text{-}prism+57aa}$, Δ*VC1807*::*P*$_{tac}$-*mNeonGreen*, Spec$^R$ | Replacement of *bap1*$^{WT}$ with a *bap1* construct lacking the prism but with remaining 57aa by cotransformation with Δ*VC1807*::*P*$_{tac}$-*mNeonGreen,* Spec$^R$ into a rugose strain lacking *rbmC* (JY071) | ZJ087 (*4*) |
| Δ*rbmC bap1*$_{Δβ\text{-}prism}$ | *vpvC*$^{W240R}$, Δ*rbmC, bap1*$_{Δβ\text{-}prism}$, Δ*VC1807*::*P*$_{tac}$-*mNeonGreen*, Spec$^R$ | Replacement of *bap1*$^{WT}$ with a *bap1* construct lacking the β-prism domain by cotransformation with Δ*VC1807*::*P*$_{tac}$-*mNeonGreen,* Spec$^R$ into a rugose strain lacking *rbmC* (JY071) | ZJ074 (*4*) |
| Δ*rbmC bap1*$_{Δ57aa*(1\text{-}repeat)}$ | *vpvC*$^{W240R}$, Δ*rbmC, bap1*$_{Δ57aa*(1\text{-}repeat)}$, Δ*VC1807*::*P*$_{tac}$-*mNeonGreen*, Spec$^R$ | Replacement of *bap1*$^{WT}$ with a *bap1* construct containing 57aa variant (1-repeat) by cotransformation with Δ*VC1807*::*P*$_{tac}$-*mNeonGreen,* Spec$^R$ into a rugose strain lacking *rbmC* (JY071) | XH108, This Study |

| | | | |
|---|---|---|---|
| Δ*rbmC* *bap1*$_{\Delta 57aa*(2\text{-}repeat)}$ | *vpvC*$^{W240R}$, Δ*rbmC*, *bap1*$_{\Delta 57aa*(2\text{-}repeat)}$, Δ*VC1807*::$P_{tac}$-*mNeonGreen*, Spec$^{R}$ | Replacement of *bap1*$^{WT}$ with a *bap1* construct containing 57aa variant (2-repeat) by cotransformation with Δ*VC1807*::$P_{tac}$-*mNeonGreen,* Spec$^{R}$ into a rugose strain lacking *rbmC* (JY071) | XH118, This Study |
| Δ*rbmC* *bap1*$_{\Delta 57aa*(core\ motif)}$ | *vpvC*$^{W240R}$, Δ*rbmC*, *bap1*$_{\Delta 57aa*(core\ motif)}$, Δ*VC1807*::$P_{tac}$-*mNeonGreen*, Spec$^{R}$ | Replacement of *bap1*$^{WT}$ with a *bap1* construct containing 57aa variant (core motif) by cotransformation with Δ*VC1807*::$P_{tac}$-*mNeonGreen,* Spec$^{R}$ into a rugose strain lacking *rbmC* (JY071) | XH194, This Study |
| Δ*rbmC* *bap1*$_{\Delta 57aa*(WFFG\text{-}>LGPE)}$ | *vpvC*$^{W240R}$, Δ*rbmC*, *bap1*$_{\Delta 57aa*(WFFG\text{-}>LGPE)}$, Δ*VC1807*::$P_{tac}$-*mNeonGreen*, Spec$^{R}$ | Replacement of *bap1*$^{WT}$ with a *bap1* construct containing 57aa variant (WFFG->LGPE) by cotransformation with Δ*VC1807*::$P_{tac}$-*mNeonGreen,* Spec$^{R}$ into a rugose strain lacking *rbmC* (JY071) | XH196, This Study |
| Δ*rbmC* *bap1*$_{\Delta 57aa+C\text{-}terminal\ 57aa}$ | *vpvC*$^{W240R}$, Δ*rbmC*, *bap1*$_{\Delta 57aa+C\text{-}terminal\ 57aa}$, Δ*VC1807*::$P_{tac}$-*mNeonGreen*, Spec$^{R}$ | Replacement of *bap1*$^{WT}$ with a *bap1* construct containing 57aa at C-terminus by cotransformation with Δ*VC1807*::$P_{tac}$-*mNeonGreen,* Spec$^{R}$ into a rugose strain lacking *rbmC* (JY071) | XH165, This Study |
| Δ*rbmC* *bap1*$_{\Delta\beta\text{-}prism+57aa*(2\text{-}repeat)}$ | *vpvC*$^{W240R}$, Δ*rbmC*, *bap1*$_{\Delta\beta\text{-}prism+57aa*(2\text{-}repeat)}$, Δ*VC1807*::$P_{tac}$-*mNeonGreen*, Spec$^{R}$ | Replacement of *bap1*$^{WT}$ with a *bap1* construct lacking the prism with remaining 57aa variant (2-repeat) by cotransformation with Δ*VC1807*::$P_{tac}$-*mNeonGreen,* Spec$^{R}$ into a rugose strain lacking *rbmC* (JY071) | XH122, This Study |
| Δ*rbmC* *bap1*$_{\Delta\beta\text{-}prism+57aa*(core\ motif)}$ | *vpvC*$^{W240R}$, Δ*rbmC*, *bap1*$_{\Delta\beta\text{-}prism+57aa*(core\ motif)}$, Δ*VC1807*::$P_{tac}$-*mNeonGreen*, Spec$^{R}$ | Replacement of *bap1*$^{WT}$ with a *bap1* construct lacking the prism with remaining 57aa variant (core motif) by cotransformation with Δ*VC1807*::$P_{tac}$-*mNeonGreen,* Spec$^{R}$ into a rugose strain lacking *rbmC* (JY071) | XH166, This Study |
| Δ*rbmC* *bap1*$_{\Delta\beta\text{-}prism+57aa*(WFFG\text{-}>LGPE)}$ | *vpvC*$^{W240R}$, Δ*rbmC*, *bap1*$_{\Delta\beta\text{-}prism+57aa*(WFFG\text{-}>LGPE)}$, Δ*VC1807*::$P_{tac}$-*mNeonGreen*, Spec$^{R}$ | Replacement of *bap1*$^{WT}$ with a *bap1* construct lacking the prism with remaining 57aa variant (WFFG->LGPE) by cotransformation with Δ*VC1807*::$P_{tac}$-*mNeonGreen,* Spec$^{R}$ | XH146, This Study |

| | | | |
|---|---|---|---|
| | | into a rugose strain lacking *rbmC* (JY071) | |
| Δ*rbmC bap1*$_{\Delta\beta\text{-prism+C-terminal 57aa}}$ | *vpvC*$^{W240R}$, Δ*rbmC, bap1*$_{\Delta\beta\text{-prism+C-terminal 57aa}}$, Δ*VC1807*::*P*$_{tac}$-*mNeonGreen*, Spec$^{R}$ | Replacement of *bap1*$^{WT}$ with a *bap1* construct lacking the prism but containing 57aa at C-terminus by cotransformation with Δ*VC1807*::*P*$_{tac}$-*mNeonGreen,* Spec$^{R}$ into a rugose strain lacking *rbmC* (JY071) | XH169, This Study |
| Δ*rbmC bap1-3XFLAG* | *vpvC*$^{W240R}$, Δ*rbmC, bap1-3XFLAG,* Δ*VC1807*::*P*$_{tac}$-*mNeonGreen,* Spec$^{R}$ | 3XFLAG tagged Bap1 with *rbmC* deleted | ZJ065 (*4*) |
| *bap1*$_{\Delta 57aa}$-*3XFLAG* | *vpvC*$^{W240R}$, *bap1*$_{\Delta 57aa}$-*3XFLAG,* Δ*VC1807*::*P*$_{tac}$-*mNeonGreen,* Spec$^{R}$ | Replacement of *bap1*$^{WT}$ with a *bap1* construct lacking the 57aa and containing a C-terminal 3XFLAG tag by cotransformation with Δ*VC1807*::*P*$_{tac}$-*mNeonGreen,* Spec$^{R}$ into a rugose strain (JY028) | ZJ036 (*4*) |
| Δ*rbmC bap1*$_{\Delta 57aa*(1\text{-repeat})}$-*3XFLAG* | *vpvC*$^{W240R}$, *bap1*$_{\Delta 57aa*(1\text{-repeat})}$-*3XFLAG,* Δ*VC1807*::*P*$_{tac}$-*mNeonGreen,* Spec$^{R}$ | Replacement of *bap1*$^{WT}$ with a *bap1* construct containing 57aa variant (1-repeat) and a C-terminal 3XFLAG tag by cotransformation with Δ*VC1807*::*P*$_{tac}$-*mNeonGreen,* Spec$^{R}$ into a rugose strain (JY028) | XH106, This Study |
| Δ*rbmC bap1*$_{\Delta 57aa*(2\text{-repeat})}$-*3XFLAG* | *vpvC*$^{W240R}$, *bap1*$_{\Delta 57aa*(2\text{-repeat})}$-*3XFLAG,* Δ*VC1807*::*P*$_{tac}$-*mNeonGreen*, Spec$^{R}$ | Replacement of *bap1*$^{WT}$ with a *bap1* construct containing 57aa variant (2-repeat) and a C-terminal 3XFLAG tag by cotransformation with Δ*VC1807*::*P*$_{tac}$-*mNeonGreen,* Spec$^{R}$ into a rugose strain (JY028) | XH198, This Study |
| Δ*rbmC bap1*$_{\Delta 57aa*(\text{core motif})}$-*3XFLAG* | *vpvC*$^{W240R}$, *bap1*$_{\Delta 57aa*(\text{core motif})}$-*3XFLAG,* Δ*VC1807*::*P*$_{tac}$-*mNeonGreen*, Spec$^{R}$ | Replacement of *bap1*$^{WT}$ with a *bap1* construct containing 57aa variant (core motif) and a C-terminal 3XFLAG tag by cotransformation with Δ*VC1807*::*P*$_{tac}$-*mNeonGreen,* Spec$^{R}$ into a rugose strain (JY028) | XH195, This Study |

| | | | |
|---|---|---|---|
| Δ*rbmC* *bap1*$_{\Delta 57aa*(WFFG\text{-}>LGPE)}$-*3XFLAG* | *vpvC*$^{W240R}$, *bap1*$_{\Delta 57aa*(WFFG\text{-}>LGPE)}$-*3XFLAG*, Δ*VC1807*::*P*$_{tac}$-*mNeonGreen*, Spec$^{R}$ | Replacement of *bap1*$^{WT}$ with a *bap1* construct containing 57aa variant (WFFG-LGPE) and a C-terminal 3XFLAG tag by cotransformation with Δ*VC1807*::*P*$_{tac}$-*mNeonGreen,* Spec$^{R}$ into a rugose strain (JY028) | XH197, This Study |
| Δ*rbmC* *bap1*$_{\Delta 57aa+C\text{-}terminal\ 57aa}$-*3XFLAG* | *vpvC*$^{W240R}$, *bap1*$_{\Delta 57aa+C\text{-}terminal\ 57aa}$-*3XFLAG*, Δ*VC1807*::*P*$_{tac}$-*mNeonGreen*, Spec$^{R}$ | Replacement of *bap1*$^{WT}$ with a *bap1* construct containing 57aa at C-terminus and a C-terminal 3XFLAG tag by cotransformation with Δ*VC1807*::*P*$_{tac}$-*mNeonGreen,* Spec$^{R}$ into a rugose strain (JY028) | XH179, This Study |
| *bap1*$_{\Delta\beta\text{-}prism+57aa}$-*3XFLAG* | *vpvC*$^{W240R}$, *bap1*$_{\Delta\beta\text{-}prism+57aa}$-*3XFLAG*, Δ*VC1807*::*P*$_{tac}$-*mNeonGreen,* Spec$^{R}$ | Replacement of *bap1*$^{WT}$ with a *bap1* construct lacking the prism but with remaining 57aa and containing a C-terminal 3XFLAG tag by cotransformation with Δ*VC1807*::*P*$_{tac}$-*mNeonGreen,* Spec$^{R}$ into a rugose strain (JY028) | XH024 (*4*) |
| Δ*rbmC bap1*$_{\Delta\beta\text{-}prism+57aa*(2\text{-}repeat)}$-*3XFLAG* | *vpvC*$^{W240R}$, *bap1*$_{\Delta\beta\text{-}prism+57aa*(2\text{-}repeat)}$-*3XFLAG*, Δ*VC1807*::*P*$_{tac}$-*mNeonGreen*, Spec$^{R}$ | Replacement of *bap1*$^{WT}$ with a *bap1* construct lacking the prism with remaining 57aa variant (2 repeats) and containing a C-terminal 3XFLAG tag by cotransformation with Δ*VC1807*::*P*$_{tac}$-*mNeonGreen,* Spec$^{R}$ into a rugose strain (JY028) | XH123, This Study |
| Δ*rbmC bap1*$_{\Delta\beta\text{-}prism+57aa*(core\ motif)}$-*3XFLAG* | *vpvC*$^{W240R}$, *bap1*$_{\Delta\beta\text{-}prism+57aa*(core\ motif)}$-*3XFLAG*, Δ*VC1807*::*P*$_{tac}$-*mNeonGreen*, Spec$^{R}$ | Replacement of *bap1*$^{WT}$ with a *bap1* construct lacking the prism with remaining 57aa variant (core motif) and containing a C-terminal 3XFLAG tag by cotransformation with Δ*VC1807*::*P*$_{tac}$-*mNeonGreen,* Spec$^{R}$ into a rugose strain (JY028) | XH174, This Study |
| Δ*rbmC bap1*$_{\Delta\beta\text{-}prism+57aa*(WFFG\text{-}>LGPE)}$-*3XFLAG* | *vpvC*$^{W240R}$, *bap1*$_{\Delta\beta\text{-}prism+57aa*(WFFG\text{-}>LGPE)}$-*3XFLAG*, Δ*VC1807*::*P*$_{tac}$-*mNeonGreen*, Spec$^{R}$ | Replacement of *bap1*$^{WT}$ with a *bap1* construct lacking the prism with remaining 57aa variant (WFFG-LGPE) and containing a C-terminal 3XFLAG tag by cotransformation with Δ*VC1807*::*P*$_{tac}$-*mNeonGreen,* Spec$^{R}$ into a rugose strain (JY028) | XH144, This Study |

| | | | |
|---|---|---|---|
| Δ*rbmC bap1*$_{\Delta\beta\text{-prism+C-terminal 57aa}}$-*3XFLAG* | *vpvC*$^{W240R}$, *bap1*$_{\Delta\beta\text{-prism+C-terminal 57aa}}$-*3XFLAG*, Δ*VC1807*::*P*$_{tac}$-*mNeonGreen*, Spec$^R$ | Replacement of *bap1*$^{WT}$ with a *bap1* construct lacking the prism but containing 57aa at C-terminus and containing a C-terminal 3XFLAG tag by cotransformation with Δ*VC1807*::*P*$_{tac}$-*mNeonGreen,* Spec$^R$ into a rugose strain (JY028) | XH184, This Study |
| Δ*rbmC bap1-3XFLAG* | *vpvC*$^{W240R}$, Δ*rbmC, bap1-3XFLAG,* Δ*VC1807*::*P*$_{tac}$-*SCFP3A,* Spec$^R$ | 3XFLAG tagged Bap1 with *rbmC* deleted | XH186, This Study |

**Table S1.**

**Strains used in this study.**

| Primer Name | Primer Sequence (5’ to 3’)* | Description |
|---|---|---|
| **Mutant constructs** | | |
| PJY140 | AATCAAACCGGGCTTTAAATTTCATCTCGAC | 3kb upstream of *bap1* |
| PJY141 | CATGATATGCAACATCTACTGAAAGAGGTGCA | 3kb downstream of *bap1* |
| PJY129 | ATATCCCGATCCAGTGCATGCAGC | 3kb upstream of *vpvc* |
| PJY130 | CCGGCTGATGCTTTGTGTCTAACGTG | 3kb downstream of *vpvc* |
| XH-U-005 | AAAACGGTTCCTTATCTAGGTGTTGAGTGGCGTACCAAAACCGTCTCTTACTCGACCACAGTACGCTATGACAT | $bap1_{\Delta 57aa*(2\text{-}repeat)}$ F |
| XH-U-006 | CCACTCAACACCTAGATAAGGAACCGTTTTAGTTTTCCACTCTAATCCTAGAGTAAACGCAGAATCTTTTGACCCC | $bap1_{\Delta 57aa*(2\text{-}repeat)}$ R |
| XH-P-041 | GGGGTCAAAAGATTCTGCGTTTACTTGGAAAACTAAAACGGTTCCTTAT | $bap1_{\Delta 57aa*(1\text{-}repeat)}$ F |
| XH-P-042 | ATGTCATAGCGTACTGTGGTCGAATAAGGAACCGTTTTAGTTTTCCA | $bap1_{\Delta 57aa*(1\text{-}repeat)}$ R |
| XH-P-045 | GTGCAACCACTGTTGATGCTCTAGGATTAGAGTGGAAAACTAAAACGGT | $bap1_{\Delta \beta\text{-}prism+57aa*(2\text{-}repeat)}$ Front F |
| XH-P-046 | ACCGTTTTAGTTTTCCACTCTAATCCTAGAGCATCAACAGTGGTTGCAC | $bap1_{\Delta \beta\text{-}prism+57aa*(2\text{-}repeat)}$ Front R |
| XH-P-047 | GGCGTACCAAAACCGTCTCTTACGTGACTGCTGACCAATCACACA | $bap1_{\Delta \beta\text{-}prism+57aa*(2\text{-}repeat)}$ Back F |
| XH-P-048 | TGTGTGATTGGTCAGCAGTCACGTAAGAGACGGTTTTGGTACGCC | $bap1_{\Delta \beta\text{-}prism+57aa*(2\text{-}repeat)}$ Back R |

| | | |
|---|---|---|
| XH-P-059 | GTACCAAAACCGTCTCTTACCTAGGCCCTGAGTGGCACACTAAACAAGTGGC | *bap1*$_{\Delta 57aa*(WFFG\text{->}LGPE)}$ F |
| XH-P-060 | GCCACTTGTTTAGTGTGCCACTCAGGGCCTAGGTAAGAGACGGTTTTGGTAC | *bap1*$_{\Delta 57aa*(WFFG\text{->}LGPE)}$ R |
| XH-P-071 | CCATTCGCGTTCCGCTGAAGTATCTAGGATTAGAGTGGAAAACTAAAACGG | Amplify 57aa for C-terminus F |
| XH-P-072 | GCGGCTGGCAGAAGTATCTTTATTTCGACAGTGTCACAGGAACG | Amplify 57aa for C-terminus R |
| XH-P-073 | CGTTCCTGTGACACTGTCGAAATAAAGATACTTCTGCCAGCCGC | 57aa insertion to C-terminus F |
| XH-P-074 | CCGTTTTAGTTTTCCACTCTAATCCTAGATACTTCAGCGGAACGCGAATGG | 57aa insertion to C-terminus R |
| XH-P-079 | TCTTACTGGTTCTTTGGCTGGCACACTAAAGTGACTGCTGACCAATCACACAT | *bap1*$_{\Delta\beta\text{-}prism+57aa*(core\ motif)}$ F |
| XH-P-080 | TTTAGTGTGCCAGCCAAAGAACCAGTAAGAAGCATCAACAGTGGTTGCACT | *bap1*$_{\Delta\beta\text{-}prism+57aa*(core\ motif)}$ R |
| XH-P-091 | aatcaccgtcatggtctttgtagtcTTTCGACAGTGTCACAGGAACG | Amplify 57aa for C-terminus FLAG R |
| XH-P-092 | CGTTCCTGTGACACTGTCGAAAgactacaaagaccatgacggtgatt | 57aa insertion to C-terminus FLAG F |
| XH-P-115 | TCTTACTGGTTCTTTGGCTGGCACACTAAATCGACCACAGTACGCTATGACAT | *bap1*$_{\Delta 57aa*(core\ motif)}$ F |
| XH-P-116 | TTTAGTGTGCCAGCCAAAGAACCAGTAAGAAGTAAACGCAGAATCTTTTGACCCC | *bap1*$_{\Delta 57aa*(core\ motif)}$ R |

| | | |
|---|---|---|
| ZJ-P-019 | CTTGTCATCGTCATCCTTGTAATCGATA | Universal sequencer for 3xFLAG |
| ZJ-P-026 | AGCAGCATTTTGAAAACTTCCGC | 2.7kb upstream of *bap1* |
| ZJ-P-027 | ATGAAATTCACGATAACCAGAAAACCG | 2.7kb downstream of *bap1* |

**Table S2.**

**Primers used in this study.**
